# Stabilisation of soil organic matter with rock dust partially counteracted by plants

**DOI:** 10.1101/2023.06.02.543347

**Authors:** Wolfram Buss, Heath Hasemer, Scott Ferguson, Justin Borevitz

## Abstract

Soil application of Ca– and Mg-rich silicates can capture and store atmospheric carbon dioxide as inorganic carbon but could also have the potential to stabilise soil organic matter (SOM). Synergies between these two processes have not been investigated. Here, we apply finely ground silicate rock mining residues (basalt and granite blend) to a loamy sand in a pot trial at a rate of 4% (equivalent to 50 t ha^-1^) and investigate the effects of a wheat plant and two watering regimes on soil carbon sequestration. Rock dust addition increased soil pH, electric conductivity and soil-exchangeable Ca and Mg contents, as expected for weathering, but decreased exchangeable levels of micronutrients Mn and Zn, likely related to soil pH. Importantly, it increased mineral-associated organic matter by 22% due to the supply of secondary minerals and associated sites for SOM sorption. Additionally, in the non-planted treatments, rock supply of Ca and Mg increased soil microaggregation that subsequently stabilised labile particulate organic matter as organic matter occluded in aggregates by 46%. Plants, however, reduced soil exchangeable Mg and Ca contents and hence counteracted the silicate rock effect on microaggregates and carbon within. We attribute this cation loss to plant exudates released to solubilise micronutrients and hence neutralise plant deficiencies. The effect of enhanced silicate rock weathering on SOM stabilisation could substantially boost its carbon sequestration potential when pH and micronutrient effects are considered.

## 1 Introduction

Urgent action is required to avoid the most dangerous impacts of climate change. Such action must include both significant reduction in greenhouse gas emissions, and atmospheric greenhouse gas removal largely via land change (IPCC, 2022). Several promising greenhouse gas removal methods are based on utilising natural cycles to capture and store atmospheric carbon dioxide (Buss, Yeates, et al., 2021; Fuss et al., 2018). These include soil organic carbon from plants and enhanced rock weathering.

During natural weathering of Ca– and Mg-rich silicates, bicarbonate (HCO_3-_) is formed, capturing CO_2_ from the atmosphere (largely as CO_2_ respired from plants and soil microorganisms) (Beerling et al., 2018; Hartmann et al., 2013). Follow-up reactions can produce solid carbonates that sequester carbon for the long-term. The mafic rock basalt is one of the most promising rock types for large-scale carbon capture and storage since it is abundant, weathers rapidly and is low in heavy metal contaminants potentially harmful for soil and plant growth. The weathering rates are highly dependent on rock particle size; 1 mm sized spheres of even the most reactive Ca– and Mg-rich silicates take thousands of years to dissolve (Hartmann et al., 2013). However, grinding rocks to a particle size of <100 µm can result in weathering and carbon drawdown on a societal relevant scale (Holdren & Speyer, 1985; Renforth, 2012). Unfortunately, across multiple studies into the weathering rates of Ca– and Mg-rich silicates in systems reflecting natural soil-plant conditions, CO_2_ sequestration rates were measured differently and varied by a factor of 1,000 (Amann et al., 2020; Haque et al., 2019, 2020; Kelland et al., 2020; ten Berge et al., 2012). This demonstrates the need for further studies that investigate enhanced weathering of globally available materials, such as mining residues, using field soils and plants but grown in controlled conditions and measured consistently across multiple interacting factors.

Soil organic carbon exists in natural systems as soil organic matter (SOM) and typically enters the soil system as plant-derived particulate organic matter (POM), which is labile and easily decomposed (Lavallee et al., 2020; Poeplau et al., 2018). Soil aggregates can physically protect POM, which is called aggregated organic matter (AggOM), and minerals can sorb partially decomposed POM fragments, so-called dissolved organic matter (DOM), to its surfaces to form mineral-associated organic matter (MAOM) (Abramoff et al., 2018; Hemingway et al., 2019; Poeplau et al., 2020). Both processes increase the retention and stability of SOM in soil. Soil aggregates are formed through various soil processes that bind together soil particles, such as activity from fungi hyphae and roots, or soil cementing agents, including Ca, Mg, Al and Fe (Amezketa, 1999). The main components of soil responsible for MAOM formation are clay and short-order Fe and Mn minerals formed from weathering of primary minerals (Kleber et al., 2015; Singh et al., 2018). Polyvalent cations, such as Ca, Mg, Al and Fe, also have a key function in facilitating sorption of DOM to mineral surfaces through cation bridging; the connection of (predominantly) negatively charged clay surfaces with negative functional groups of DOM (Kleber et al., 2015; Singh et al., 2018).

Synergies between enhanced rock weathering for both the formation of inorganic carbon for direct carbon drawdown and the formation of secondary minerals and polyvalent cations for stabilisation of SOM could significantly enhance its carbon sequestration potential and have soil health co-benefits. Yet studies in this area that investigate the factors that influence its potential are lacking.

Both rock weathering rates and SOM formation and decomposition reactions are affected by soil water availability and plant activity. Higher precipitation and increased water flow accelerates rock weathering (Brady et al., 1999; G. Li et al., 2016; White & Blum, 1995) and water availability also governs microbial processes responsible for SOM decomposition and affects plants that supply carbon into soil. Therefore, precipitation has a strong effect on SOM levels (Luo et al., 2017) and the SOM content is typically higher in areas with more precipitation (Alvarez, 2005; Wiesmeier et al., 2019). Plants can significantly increase rock weathering rates by up 10-fold (Bormann et al., 1998; Cochran & Berner, 1996; Hinsinger et al., 2001). Plants also provide the foundation for SOM formation through rhizodeposits from living plants and litter from dead plants. Yet, they can also accelerate the decomposition of existing SOM through various processes summarised under the term positive priming (Keiluweit et al., 2015; Kuzyakov, 2010). The effects of plants and water could significantly influence the potential synergies of rock weathering on SOM and inorganic carbon sequestration.

In this study, to simulate potential field conditions, bulk mining residues (basalt, granite blend) were applied to a sandy loam and field climatic conditions were replicated in a growth chamber under controlled conditions. The effects of wheat plants and watering on rock weathering, microbial composition and SOM content of different stability were investigated. Further, the soil available and plant tissue elemental contents were analysed to understand the mechanisms behind the soil response. The hypothesis was that weathering of Ca– and Mgrich silicates can increase both inorganic and organic carbon storage.

## 2 Materials and Methods

### 2.1 Soil and rock samples

The soil was agricultural topsoil (0-20 cm) sourced in 2020 from Young in central New South Wales, Australia. The soil was dried and stored for ∼12 months before it was used in the incubation trial. It had a pH (in water) of 5.68 and was classified as loamy sand (USDA classification). The cation exchange capacity was 2.3 cmol_+_ kg^-1^ and the total carbon and nitrogen contents were 0.88% and 0.045%, respectively. More details about the soil with full characterisation can be found in SI Table 1.

**Table 1:** Characterisation of rock sample.

|  |  |
| --- | --- |
| Mineralogy | (%) |
| Amorphous | 19.1 |
| Quartz | 16.3 |
| Plagioclase | 34.4 |
| K-feldspar | 5.7 |
| Clinopyroxene | 10.4 |
| Olivine | 2 |
| Nepheline | 1.2 |
| Analcime | 1.3 |
| Apatite | 1.3 |
| Kaolinite | 0.2 |
| Mica | 4.4 |
| 2:1 Clay | 3.7 |
| * Chlorite, Vermiculite or Smectite |  |
| Elemental content | (%) |
| Al<sub>2</sub>O<sub>3</sub> | 13.4 |
| BaO | 0.04 |
| CaO | 4.67 |
| Cr<sub>2</sub>O<sub>3</sub> | 0.05 |
| Fe<sub>2</sub>O<sub>3</sub> | 8.12 |
| K<sub>2</sub>O | 2.65 |
| MgO | 4.13 |
| MnO | 0.10 |
| Na<sub>2</sub>O | 3.24 |
| P<sub>2</sub>O<sub>5</sub> | 0.42 |
| SiO<sub>2</sub> | 60.9 |
| SrO | 0.05 |
| TiO<sub>2</sub> | 1.41 |
| Particle size | (%) |
| <90 | 96.5 |
| <70 | 89.3 |
| <50 | 80.1 |
| <10 | 29.6 |
| Loss on ignition | 0.68 |
| pH (in water) | 9.2 |
| C content (%) | 0.075 |

The rock mining residues were sourced from Victoria in Australia (Cohuna and Carisbrook) and comprise of both basalt and granite. The rock was ground with a pug mill and sieved to <90 µm with a p80 (80% of particles with a diameter less than specified size) of 50 µm. Data on X-ray diffraction and full acid digestion followed by inductively coupled plasma (ICP) – mass spectrometry (method ME-MS61 using perchloric, nitric, hydrofluoric and hydrochloric acids) and particle size using a Mastersizer 2000 (Malvern Panalytical; Malvern, UK) are shown in Table 1 and full particle size distribution of both materials in SI Figure 1.

**Figure 1:**
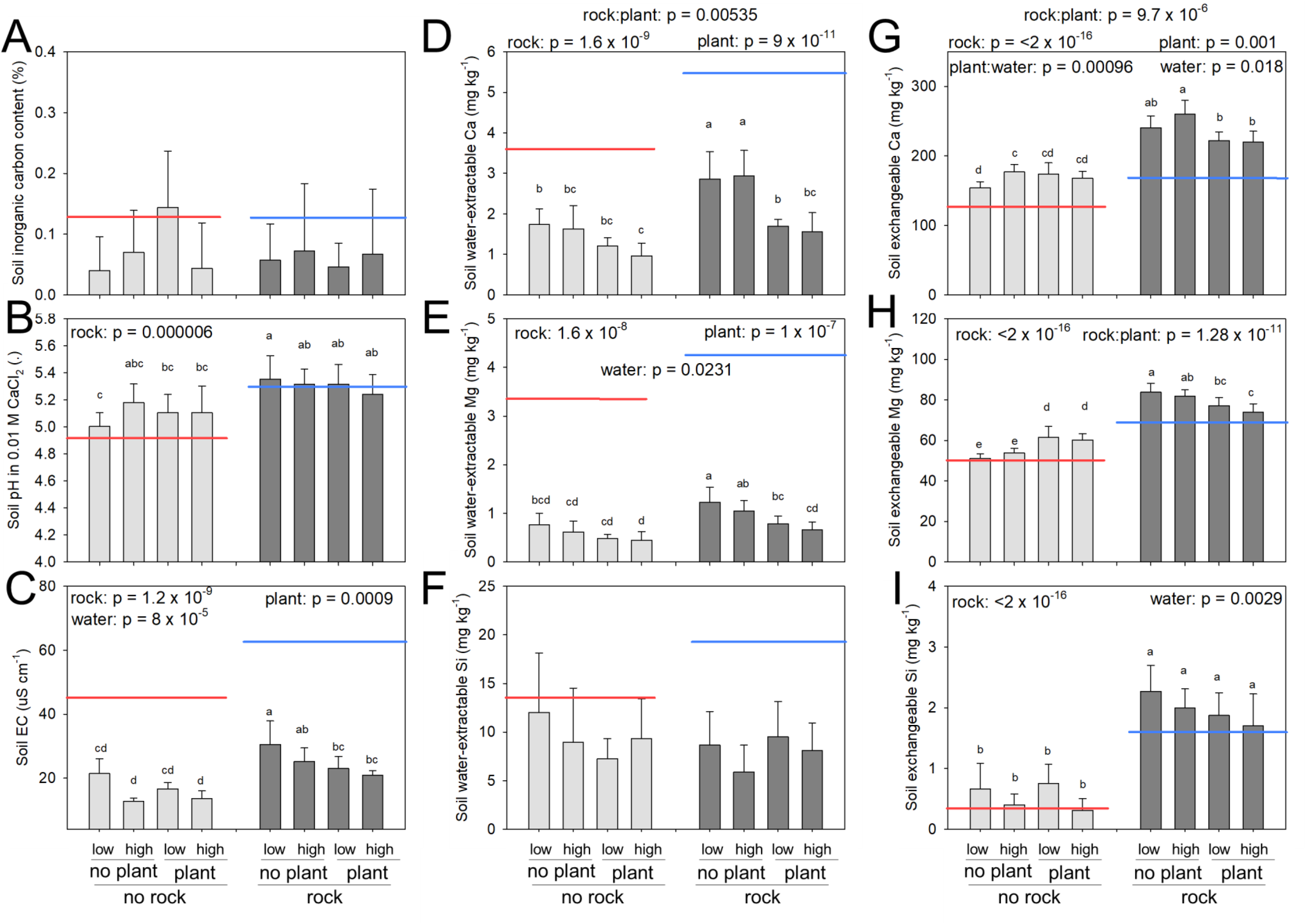
Soil indicators for weathering of Ca– and Mg-rich silicate rocks at the end of 6-month incubation. (A) Inorganic carbon content, (B) pH, (C) EC, and (D) water-extractable Ca, (E) Mg and (F) Si contents and ammonium acetate-extractable (exchangeable) (G) Ca, (H) Mg and (I) Si contents. Soil was analysed at the end of a 6-month incubation study using a full factorial design with high/low water, unplanted/planted soil and soil only/rock addition. Red and blue lines indicate baseline values from the soil at the start of the trial without and with rock addition (∼4%), respectively. Main effects determined using one-way ANOVAs (results shown at the top of each figure). Different letters indicate significant differences among the treatments determined via Tukey post-hoc test.

### 2.2 X-ray diffraction of rock sample

Samples were pre-ground and sieved to <90 µm, then spiked with 20 wt% Al_2_O_3_ (Baikalox polishing corundum) and manually ground finely in an agate mortar in acetone. The suspension was pipetted on low-background holders (quartz), and dried. Powder X-ray diffraction analysis was carried out with a Malvern Panalytical Empyrean Series 3 diffractometer that was equipped with Bragg-Brentano^HD^ divergent beam optic and a PIXcel^3D^ detector (1D scanning mode, 3.347° active length), using Co*Ka* radiation. Samples were analysed over a range of 4-85° 2*q*, with step width of 0.0131303° 2*q* and a total dwell time of 98 s/step, while spinning samples horizontally. Phase identification was carried out with the software Diffrac*Plus* Eva 10 (2004; Bruker AXS GmbH, Karlsruhe, Germany) and ICDD PDF-2 database (2004; PDF-2. International Centre for Diffraction Data, Newtown Square, PA, USA), and quantification with Siroquant V4 (Taylor, 1991) and Highscore Plus 4.8 (2018; Malvern Panalytical B. V., Almelo, The Netherlands).

### 2.3 Soil incubation / wheat plant trial

The soil was sieved to <10 mm to remove large root and other plant structures. Soil or crushed rock-soil mix (1.2 kg total) was filled into round pots 11 cm diameter and 11 cm high with drain holes. Crushed rock was pre-mixed with the soil in ziplock bags at a rate of ∼4% w/w (equivalent to 50 t ha^-1^). After watering, pre-germinated wheat seedlings (*Triticum aestivum*, Condo variety) were planted into the centre of the pots (half of the pots were planted).

A full-factorial design with treatments of rock/no rock, wheat/no wheat and low/high water was used with 8 pot replicates (64 pots in total). No fertiliser was applied, and the pots were kept in a growth chamber in a randomised block design for ∼6 months simulating the diurnal and seasonal light and temperature regime of central New South Wales, Australia, the origin of the soil (temperature profile of ∼8-10°C at night and ∼18-25°C during the day). Pots were watered with pre-determined amounts of tap water depending on high and low water treatments. The high-water treatment received 100 mL of water three times a week and the low water treatment 100 mL and 50 mL water each week. In some weeks lower/high watering was necessary depending on plant growth stage resulting in 7 L and 4 L of water in total for the high– and low-water treatments, which corresponds to ∼740 and 420 mm precipitation over the 6 months. This is approximately equivalent to the annual lower and higher end of the precipitation in the area the soil and light/temperature regime were adapted from.

At the end of the ∼6 month-period, the plants and their roots were pulled out of the soil and partitioned into roots, shoots and head (seeds/grain), dried at 105°C and weighed. Centrifuge tubes (50 mL) were used to take soil samples from the area under the plant at harvest (3 cm deep core). The soil samples were dried in the oven at 40°C for 3 days.

### 2.4 Soil analysis

#### 2.4.1 Carbon analysis

A soil subsample (∼400 mg) was ground, and the contents of carbon and nitrogen were determined with a VarioMax 3000 (Elementar, Germany) (peak anticipated N: 210 s, oxygen dosing time: 15 s, oxygen dosing: 70 ml min^-1^, furnace temperatures: 900°C, 900°C and 830°C; helium as carrier gas). The inorganic carbon content was determined through prior soil acidification using 1 M HCl until no gas release was visible.

#### 2.4.2 Three-pool soil carbon fractionation

Full details about the soil carbon fractionation that distinguishes free POM, AggOM and MAOM is described previously (Buss, Sharma, et al., 2021). In short: 10 g of soil was shaken with a total volume of 50 mL of deionised water. Wet sieving was performed using 70 µm sieves. For separating POM and AggOM sodium iodide adjusted to a density of 1.8 g cm^-3^ was used. ^33^The samples were centrifuged at 3000 rpm for 10 min and POM and AggOM were separated by decanting the content of the tube onto a Whatman No. 2 filter paper. To remove sodium iodide residues, the AggOM fraction was washed with deionised water. The MAOM fraction (derived from the sieving step; <70 µm) was separated from the liquid fraction via centrifugation at 3000 rpm for 30 min.

The carbon content within each fraction was analysed using a combustion method (details above). The aqueous fraction was analysed for electric conductivity (EC) with a ProLab 5000 pH/EC meter (SI Analytics; Germany) and a suite of elements using ICP (details about ICP below). The elemental content determined in this fraction is referred to as “water-extractable content” in the following.

#### 2.4.3 Two-pool soil carbon fractionation

The AggOM fraction comprises both occluded POM, protected from decomposition, but also aggregated clay and silt particles that contain sorbed carbon (MAOM), sourced from either plant exudates or residual POM decomposition (SI Figure 2). To further separate AggOM into occluded (and free) POM and MAOM, so to investigate whether indeed carbon was occluded in aggregates or whether it was only sorbed to the extra mineral surfaces provided by the weathered rock, a second fractionation based on previous work (Cotrufo et al., 2019) with some modifications was applied on a subset of samples (no rock-no plant; rock-no plant; rock-plant).

**Figure 2:**
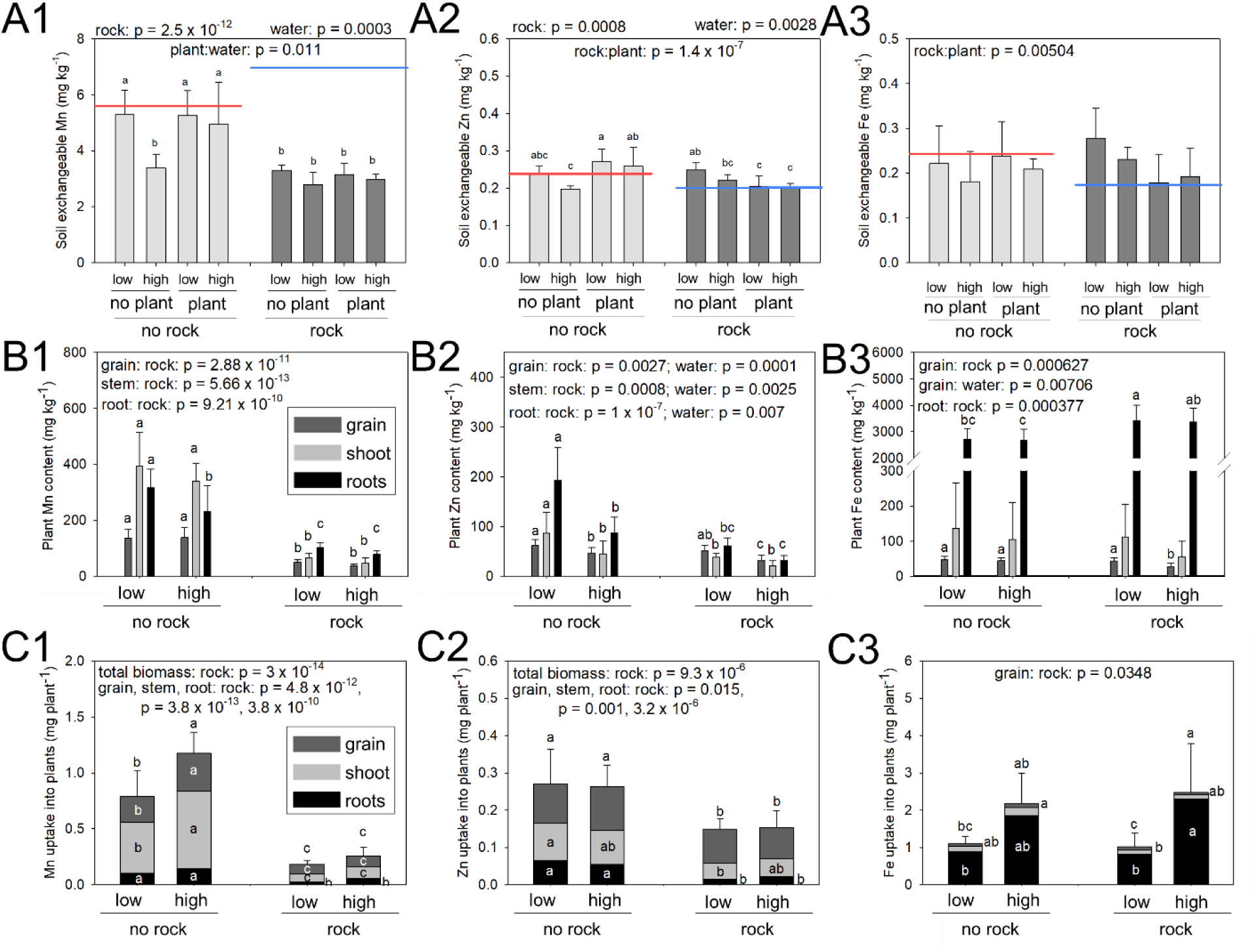
Micronutrients Mn (1), Zn (2) and (3) Fe in plant and soil at the end of 6-month incubation. (A) Ammonium acetate exchangeable contents for soils in all 8 treatments, and (B) plant tissue contents of grain, shoots and roots and (C) total plant uptake in grain, shoot and roots for the planted treatments. Red and blue lines indicate baseline values from the soil at the start of the trial without and with rock addition (∼4%), respectively. Main effects determined using one-way ANOVAs (results shown at the top of each figure). Different letters indicate significant differences among the treatments determined via Tukey post-hoc test.

A solution with 0.5% hexameta phosphate was prepared, and 30 mL added to 50 mL centrifuge tubes that contained 2.5 g of soil and 2 glass beads. The tubes were shaken at 150 rpm for 18 hours and then sieved through 70 µm sieves (free and occluded POM). The MAOM fraction was subsequently separated from the aqueous fraction through centrifugation at 3000 rpm for 30 min. The free and occluded POM, MAOM and aqueous fraction were all analysed for their carbon content.

#### 2.4.4 ICP-Optical Emission Spectroscopy (ICP-OES)

The aqueous fractions recovered from the fractionation described under 2.3.2 were analysed with an ICP-OES 5110 (Agilent; Santa Clara, CA, USA) for 20 elements. The ICP multi-element standard solution Intelliquant No.1 and 2 from Agilent were used for calibration using following concentrations: calibration blank, 0.01, 0.05, 0.1, 0.01, 1, 5, 10 and 50 mg L-^1^. The 1 ppm standard was used as internal quality control.

#### 2.4.5 Soil aggregation test

Soil aggregation was tested on a sub-set of samples (no rock-no plant; rock-no plant; rock-plant) to investigate which aggregates where increased in the rock-no plant treatment that resulted in the elevated AggOM contents. To determine soil aggregates, 7 sieves (sizes: 1000, 500, 400, 300, 200 and 70 µm; pluriStrainer, pluriSelect, Leipzig, Germany) were stacked on 50 mL centrifuge tubes. Suction was applied with a syringe to facilitate filtering of 1 g of soil and 500 mL of water through the sieves. Subsequently, the sieves were dried and weighed.

The amount of soil in the <70 µm fraction was determined by the difference in mass.

#### 2.4.6 Soil pH measurement

The soil pH was measured by shaking 1.5 g of soil with 30 mL of either deionised water or 0.01 M CaCl_2_ in 50 mL tubes at 150 rpm for 1 hour. Tubes were left to settle for 20 min before the pH was measured with a ProLab 5000 pH/EC meter (SI Analytics; Germany).

### 2.5 Plant elemental content

Representative samples of grain, stem and roots (250 mg) were digested in 9 mL concentrated HNO_3_ and 1 mL H_2_O_2_ in a microwave digester (Milestone ETHOS UP) at 210°C and 1800 W for 35 min. Samples were subsequently diluted to 2% HNO_3_ and analysed via ICP-OES (as described above).

### 2.6 Microbial composition via shotgun metagenomics

Full details about the method to determine microbial composition was described previously (Buss, Sharma, et al., 2021). In short: a commercial DNA extraction kit (DNeasy PowerSoil Pro Kit, Qiagen, Hilden, Germany) was used for extracting DNA from the dried soil and the samples were barcoded with the “Native Barcoding kit” (Oxford Nanopore Technology, Oxford, UK) and run on a Flongle flow cell (Oxford Nanopore Technology, Oxford, UK).

Data were basecalled and demultiplexed with Guppy (version: 5.0.7; Oxford Nanopore Technology, Oxford, UK and all short (<200 bp) and low quality (<q7) sequences were removed with NanoPack (De Coster et al., 2018).

Sequences were blasted against the NCBI nucleotide database version 5 (Sayers et al., 2021). The taxonomic ID for the single best blast hit per sequence was extracted. Sequences without a match and operational taxonomic units (phylum, class or order) that were only detected once in a sample were excluded. Sequences matching an operational taxonomic unit (phylum, class or order) observed in less than 8 samples (each treatment had 8 replicates) were also filtered out.

### 2.7 Statistics

Statistical analyses were conducted in R (2022.12.0) and most visualisations in SigmaPlot (Systat Software Inc; San Jose CA, USA). Analysis of variance and Tukey post-hoc tests were conducted in R using the aov and TukeyHSD functions.

Microbial data were processed with Analysis of compositions of microbiomes with bias correction (ANCOM-BC) on phyla, class and order level using the ANCOMBC2 package in R (H. Lin & Peddada, 2020). The method normalises the data and checks for statistical differences based on treatment effects. In addition, the row microbial count data (on phylum, class and order level) were used for clustering using Non-Metric Multi-Dimensional Scaling (NMDS) (vegan package; metaMDS; Bray-Curtis dissimilarity matrix) and NMDS 1 and 2 were subsequently plotted. PERMANOVA (vegan package in R) was used to check for significant differences as a result of rock, plant and water treatments.

## 3 Results

### 3.1 Rock weathering, soil minerals and plant uptake

The inorganic carbon was not different between the control and rock amended treatments (Figure 1A). However, rock dust addition increased soil pH and EC immediately after application to soil (blue – rock dust baseline in Figure 1B and C) and the rock effect persisted throughout the incubation (significant rock effect on pH and EC). EC decreased from the baseline values in both the soil only and rock amended treatments. Plants and a high-water treatment accelerated EC decrease (Figure 1C).

Water-extractable Ca and Mg contents in soil increased significantly after rock addition as expected for weathering, but the plant effect eliminated this and decreased water-extractable Ca and Mg down to values comparable to the no-rock control treatments (Figure 1D, E).

Rock addition also increased the ammonium acetate-extractable (exchangeable) Ca and Mg contents, and the levels were significantly higher after the incubation for both planted and unplanted treatments (Figure 1G and H). There was an initial peak of Ca and Mg release of fresh rock-sand samples, but we could not detect associated changes with any bi-(carbonate) levels in water-extractions (data not shown), predicted as part of alkalinity release during rock weathering. In contrast to the water-extractable contents of Ca and Mg that decreased over the course of the incubation compared to the baseline value (blue line in Figure 1D, E), the ammonium acetate-exchangeable content increased (Figure 1G, H). The increase in soil exchangeable Ca is highly significant in all rock treatments (p-values < 0.0001). For Mg only the two unplanted treatments were significantly higher than the rock-sand baseline (low water: p = 0.00003; high water: p = 0.00029). The water-extractable Si content did not change as a result of rock addition (Figure 1F), but the exchangeable content increased by 3-4-fold (Figure 1I).

There was no overall effect of rock addition on plant biomass or the individual plant parts (SI Figure 3) and no effect on uptake (total mass) of Ca and Mg into plant tissue (SI Figure 4).

**Figure 3:**
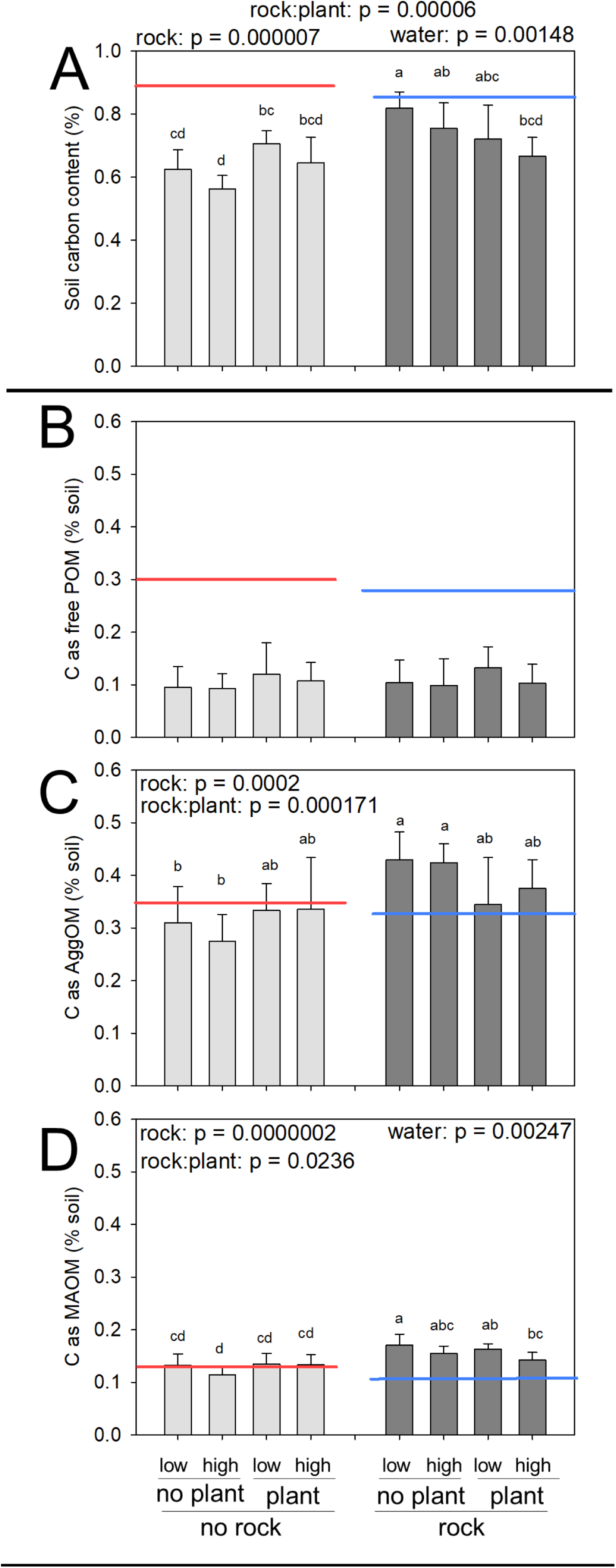
(A) Total soil carbon content and (B-D) three-pool soil carbon fractionation data at the end of 6-month incubation. (B-D) show carbon associated with (B) particulate organic matter (POM), (C) aggregate organic matter (AggOM) and (D) mineral-associated organic matter (MAOM). A full factorial design of high/low water, unplanted/planted soil and soil only/rock addition was used. Red and blue lines indicate baseline values from the soil at the start of the trial without and with rock addition (∼4%), respectively. Main effects determined using one-way ANOVAs (results shown at the top of each figure). Different letters indicate significant differences among the treatments determined via Tukey post-hoc test.

**Figure 4:**
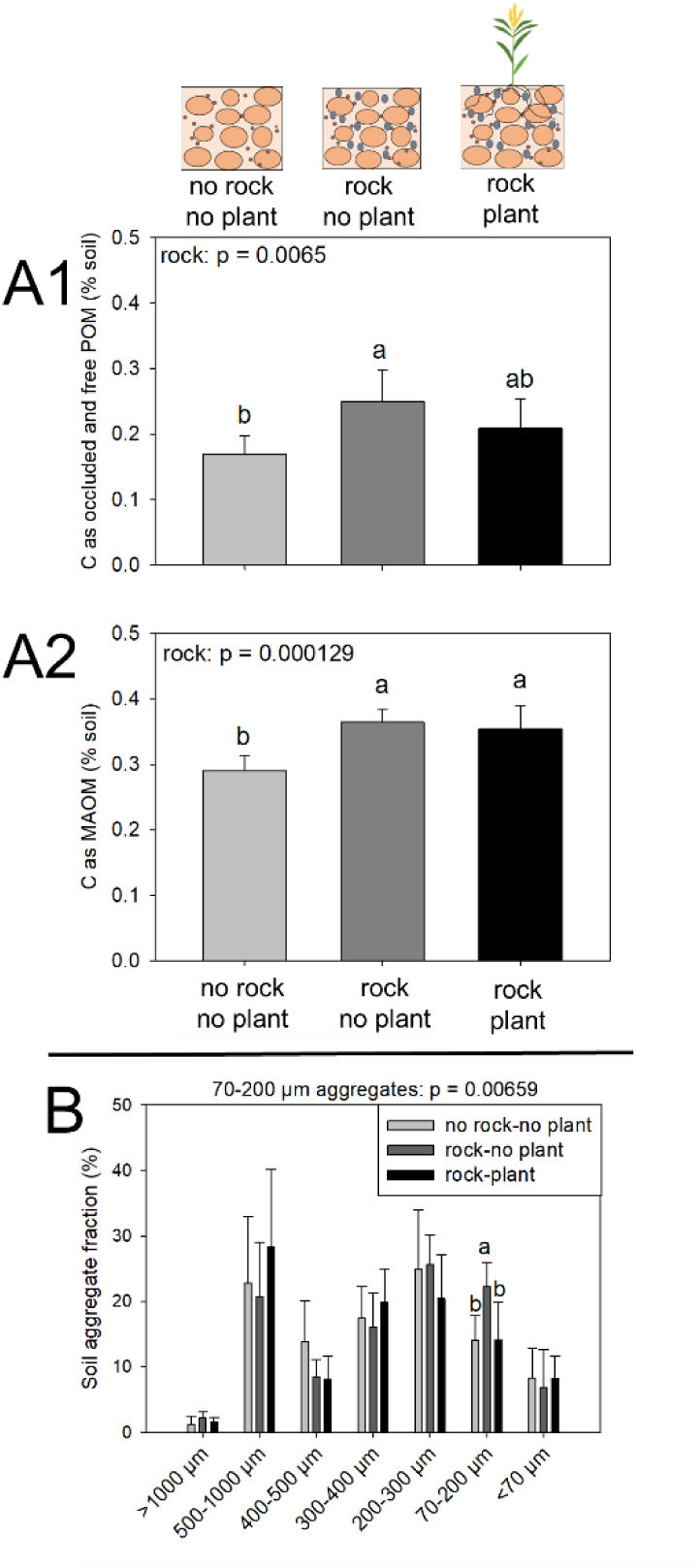
(A) Soil carbon occlusion determined via 2-pool soil fractionation and (B) soil aggregates in a subset of samples. Treatments: no rock without plant (control-control-low), rock with and without plants (all low water treatments). (A) Soil carbon fractionation using hexameta-phosphate extraction for full soil disaggregation at the end of a 6-months incubation. Carbon associated with the (A1) free and occluded POM fraction and (A2) MAOM fraction. Main effects determined using one-way ANOVAs (results shown at the top of each figure). Different letters indicate significant differences among the treatments determined via Tukey post-hoc test. (B) Soil aggregate classes. For statistical analysis (ANOVA, followed by Tukey’s post-hoc test), data were centred log ratio transformed.

But the plant grain Ca content (mg Ca per kg plant biomass) and total Ca uptake into grain (mg Ca per plant) significantly increased because of rock addition (p = 3.9 10^-6^; SI Figure 4 and SI Figure 5). Plant tissue contents (stem and root) and total uptake of Si in plant tissue increased significantly due to rock application (SI Figure 4 and SI Figure 5).

**Figure 5:**
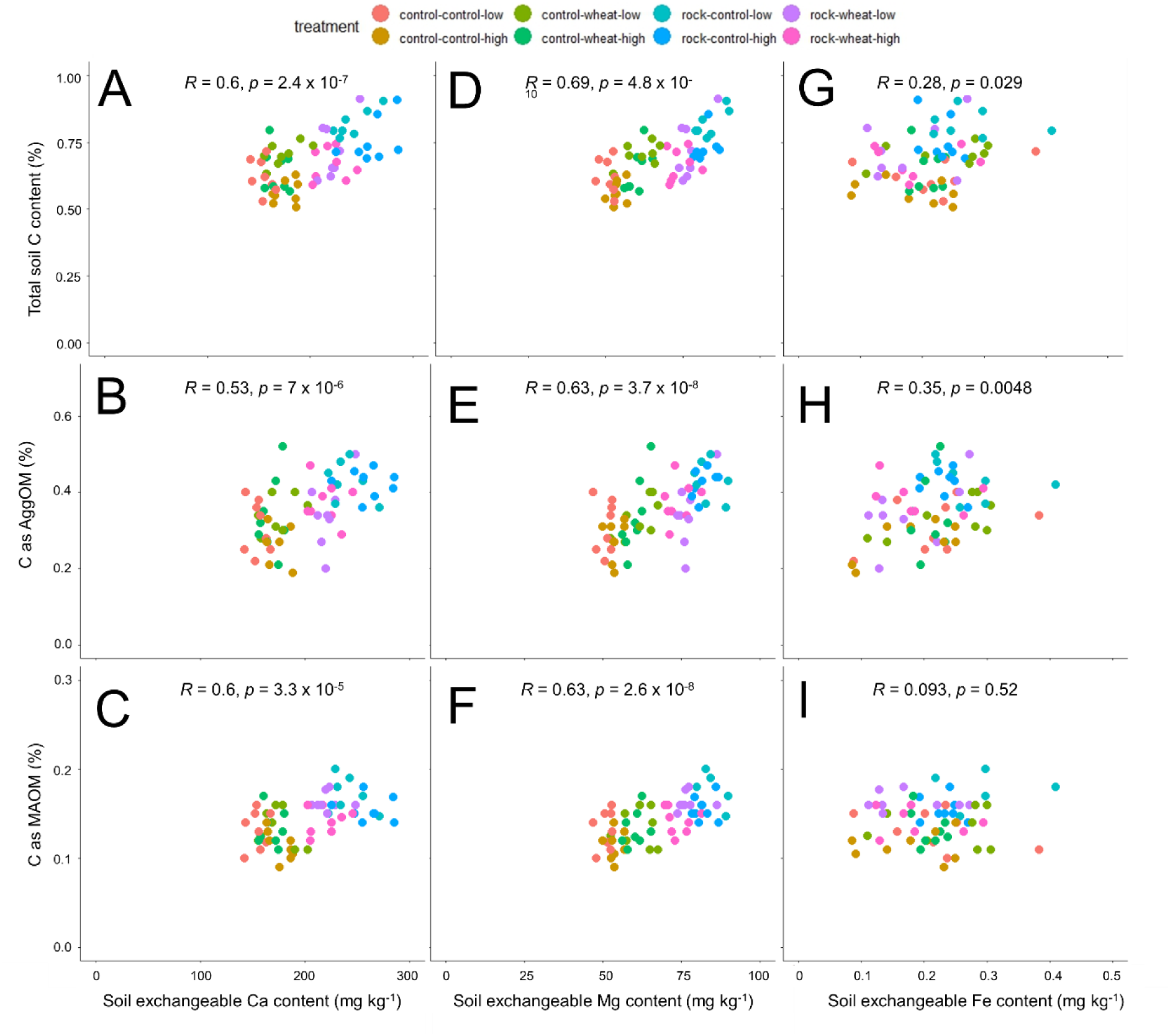
Relationship of soil exchangeable (A-C) Ca, (D-F) Mg and (G-H) Fe contents with (A, D, G) total soil C content, (B, E, H) C associated with AggOM and (C, F, I) C associated with MAOM at the end of 6-months incubation. Treatment labels are control/rock – control/wheat – low/high water. Pearson correlation coefficient and p-value shown, respectively.

Rock amendment significantly decreased soil exchangeable contents, plant tissue levels and plant uptake of micronutrients Mn and Zn (Figure 2). The rock treatment did not significantly reduce exchangeable Fe levels (Figure 2A3), yet within the rock treatments, plant addition decreased exchangeable Fe (Figure 2A3) and Zn (Figure 2A2). Rock significantly decreased the content and total uptake of Fe into grains (Figure 2C3).

### 3.2 Soil carbon content and three-pool soil carbon fractionation

The total soil carbon content decreased over the course of the incubation in all treatments (red/blue lines in Figure 3A). The rock-amended soils started with a slightly lower soil carbon content (blue line) than the control (red line) because the mass of rock addition diluted the soil carbon content (rock 0.075% C; soil 0.88% C).

By the end of the experiment, rock addition led to a 16% higher total carbon content across all treatments (rock effect: p = 0.00007; Figure 3A) and an even 32% higher content when only the non-planted treatments are considered. Plants partially counteracted the rock effect decreasing soil carbon content (rock:plant effect: p = 0.00006). Overall, the three treatments and their interactions explained 56% of the variance in soil carbon content of which 35% was explained by rock, 5% by water, 9% by plants and 19% by rock:plant interactions.

We next conducted a three-pool soil fractions to separate labile (free) POM, mineral and aggregate components (SI Figure 2). The carbon content associated with the free POM fraction decreased drastically over the course of the trial in all treatments from ∼0.3% in the baseline (red/blue line) to ∼0.1% (Figure 3B). None of the treatments affected the free POM loss. Rock addition significantly increased C associated with AggOM by 25% over the non-amended control and by a massive 46% when only the non-planted treatments are considered (Figure 3C). Plants counteracted the effect of rock-amendment on AggOM (rock:plant effect; p = 0.0002). The carbon content associated with the MAOM fraction increased by 22% because of rock addition and by 32% when only the non-planted treatments are considered (Figure 3D). Rock-amendment significantly increased the amount of soil recovered as MAOM (SI Figure 6A) but decreased the concentration of carbon within the MAOM fraction by 14% (p = 0.0002; SI Figure 6B).

**Figure 6:**
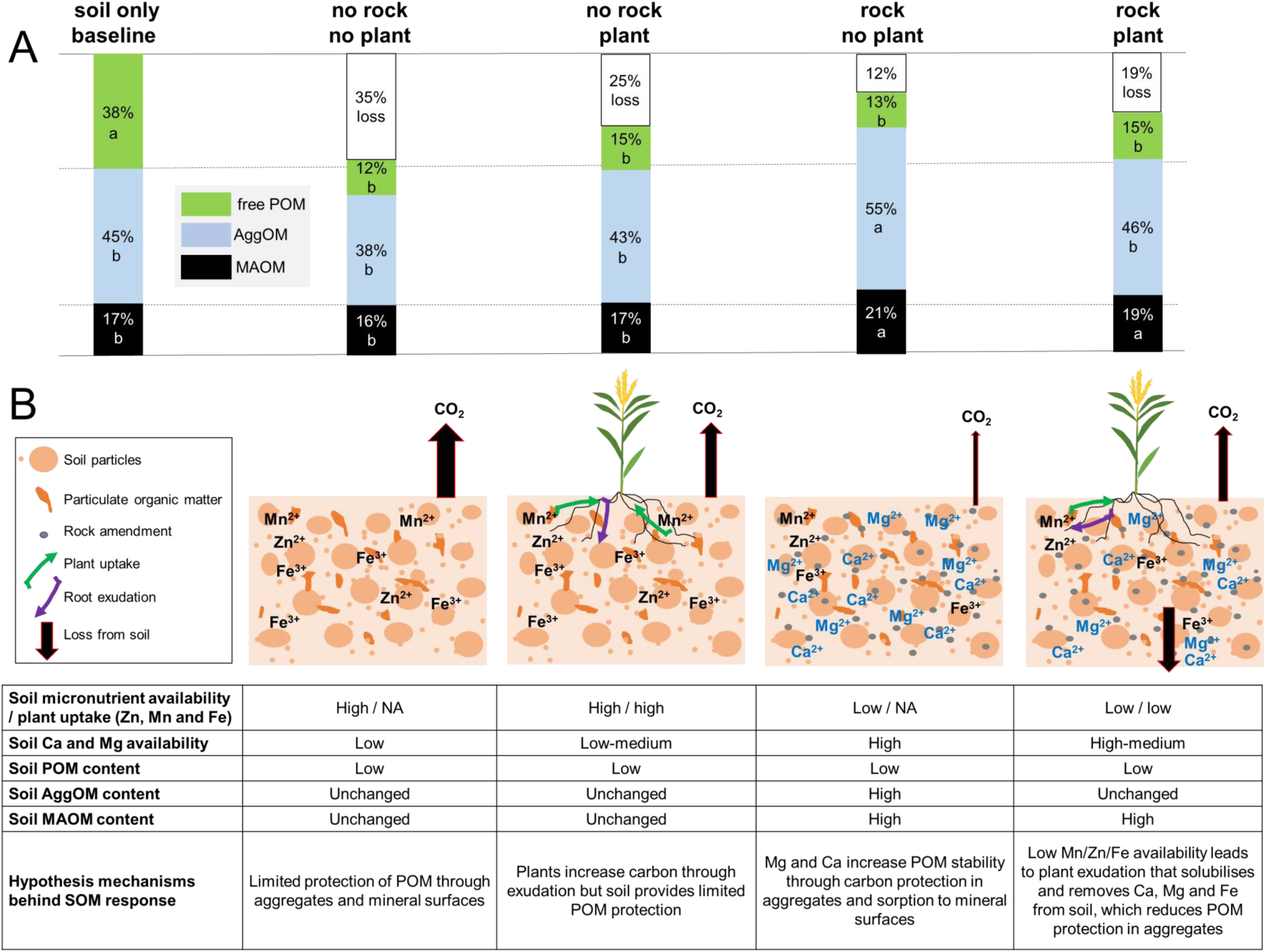
Summary of soil (carbon) responses to rock and plant additions relative to the soil baseline. (A) Conversion of SOM fractions in the baseline scenario (soil before trial) into other soil carbon fractions based on data in Figure 3 (low– and high-water treatments pooled together, n = 16). Letters show significant differences among the treatments. (B) Schematic summary of the key results.

High water treatment resulted in significantly lower soil carbon levels than low water treatment (p = 0.00148; Figure 3A). Water treatment only affected the carbon content associated with MAOM fraction (Figure 3D; p = 0.002). This loss of carbon was not associated with a lower carbon content within the MAOM fraction (SI Figure 6B), but instead with less soil recovered in this fraction (SI Figure 6A; water effect: p = 0.0006).

### 3.3 Two-pool soil carbon fractionation and soil aggregates on a subset of samples

Using a two-pool soil carbon fractionation technique that ensures full disaggregation (and hence separation of the AggOM pool into MAOM and free plus occluded POM) on a sub-set of samples (Figure 4A; schematic SI Figure 2), both planted and unplanted rock treatments significantly increased the carbon content associated with the MAOM fraction by ∼0.07% compared to the no-rock treatment. Rock addition also increased the POM fraction that includes both occluded and free POM (Figure 4A1; p = 0.006), but only the unplanted, rock-amended treatment was different to the control (p = 0.005; Figure 4A1). The aqueous fraction that is used to extract the soil, contained dissolved organic matter (DOM) in the range of 0.12-0.15% carbon per unit of soil (SI Figure 7). There was a statistically significant increase in the DOM pool in the rock, non-planted treatment compared to the control treatment (ANOVA: p = 0.049; Tukey: p = 0.046).

In the same subset of samples, we analysed soil aggregation within aggregate size classes of <70 µm to >1000 µm (Figure 4B). There was a statistically significant increase in microaggregates of size 70-200 µm from 14.1% in the control to 22.2% in the unplanted, rock-amended (Tukey post-hoc test: p = 0.004). Plants reduced the percentage of microaggregates to 14.2%, fully counteracting the increase in microaggregation induced by rock addition (Figure 4B).

### 3.4 Soil DNA and microbial composition

The extractable soil DNA content was significantly higher in rock-amended treatments compared to unamended treatments (SI Figure 8A). The DNA content decreased over the course of the trial compared to the baseline indicating that soil DNA was lost. There was a significant correlation between soil DNA and soil carbon content (SI Figure 8B).

The total number of continuous DNA fragments (reads) extracted and sequenced per sample were between 4,000 to 50,000 with an average read length of 600-3,000 base pairs (SI Table 3), generating >1Gb of sequence. The abundant microbial high level taxonomic groups detected via shotgun metagenomics and long-read sequencing did not change significantly as a result of rock addition as shown via clustering (SI Figure 9B) and percentage composition (SI Figure 10). We also did not detect specific changes due to basalt or plant effects on individual microbial taxa (SI Table 4). Water treatment, however, significantly affected clustering of the samples based on phylum, class and order level (phylum level clustering in SI Figure 9A (PERMANOVA results in SI Table 5). There were also significant effects within the microbial taxa due to water treatment at the class and order level (SI Table 4).

### 3.5 Associations of soil properties with SOM fractions and SOM transformations

There was a highly significant correlation between soil exchangeable Ca and Mg contents and soil carbon content (p = 2.4 x 10^-7^ and p = 2.4 x 10^-10^; Figure 5A,D) and a significant correlation with Fe content (p = 0.029; Figure 5G). Exchangeable Ca and Mg contents also correlated highly significantly with both carbon in AggOM (Figure 5B and E) and MAOM (Figure 5C and F). Exchangeable Fe only correlated highly significantly with carbon as AggOM (p = 0.005; Figure 5H).

The schematic in Figure 6A shows the effects of rock and rock-plant interactions on soil carbon of different stability compared to the soil-only baseline and is based on Figure 3B-D. Low– and high-water treatments were pooled to focus on plant and rock effects. C associated with POM was reduced in all treatments compared to the baseline. However, depending on treatment, the carbon was either lost or converted into different soil carbon fractions (Figure 6A). In the no-rock treatments, free POM was lost without any conversion into AggOM or MAOM (C content as AggOM and MAOM same as in the baseline). In the rock amended treatment, some free POM was instead retained as MAOM and as AggOM. In the non– planted, rock amended treatment 12% of POM was lost and 19% in the treatment with plant. Hence, the rock-amended, non-planted control retained 23% of free POM lost in the control sample in the form of more stable soil carbon fractions.

## 4 Discussion

### 4.1 Inorganic carbon sequestration

Previous rock weathering studies have found it is challenging to directly detect changes to soil inorganic carbon content and therefore, typically proxies are used to assess rock weathering and associated drawdown of atmospheric CO_2_. Such proxies include Ca and Mg mass balance approaches based on both pore water or ammonium acetate extractable cations and changes in pH and EC compared to a non-amended control (Amann et al., 2020; Amann & Hartmann, 2022; Kelland et al., 2020; ten Berge et al., 2012). In this study, we found elevated levels of water-extractable and exchangeable Ca and Mg contents and pH and EC levels in soil 6 months after rock addition. However, water-extractable Ca and Mg did not indicate additional weathering of our mining residues since the levels peaked at the start of the incubation (fresh rock-soil mix) and this peak was not associated with any (bi)carbonate formation (product of Ca– and Mg-rich silicate reaction with carbonic acid). Our mining residues had pre-weathered and contained 19.1% amorphous material and 3.9% secondary minerals, which likely contained cations that were only sorbed to mineral surfaces and were released without reacting with carbonic acid. These readily available cations would be responsible for this initial Ca and Mg release. Pre-weathering eliminates the Ca and Mg mass balance approach as proxy for new carbon drawdown and may be unsuitable for mining residues, at least if water extraction or pore water values are used.

There is some evidence for rock weathering based on exchangeable Ca and Mg contents, which were higher at the end of the trial compared to the rock-soil baseline. However, there is also no (extra) inorganic carbon detected in soil at the end of the incubation, as previously reported (Kelland et al., 2020). Overall, we could not find evidence for inorganic carbon formation in our trial. Only around half of the rock was composed of basalt and the percentage of the fast-weathering mineral olivine was only 2%, which expectantly did not result in a strong weathering signal at an application rate of 50 t ha^-1^. Given the large range of weathering rates and associated carbon drawdown rates in the literature, it is clear that the method for determining weathering rates needs refinement. Other indirect weathering effects, as presented here for SOM, do show potential.

### 4.2 Rock addition increases organic carbon protection

POM in sandy soil has little protection from decomposition and subsequently 2/3 of the POM in the baseline soil was lost after 6 months of incubation under growing conditions (Figure 6A). Rock dust addition decreased SOM losses by transformation of POM into more durable soil carbon fractions, and the effects were linked to changes in soil chemistry. We failed to detect substantial abundance or composition shifts in the microbiome as a result of rock addition (SI Figure 9). Our microbiome assay using shotgun, long-read sequences via Oxford Nanopore Technology is robust and can detect differences among treatments in environmental samples (Hamner et al., 2019; Loit et al., 2019; Petersen et al., 2020). Our study shows that water treatment did have an effect on microbial composition as expected. The soil response to rock addition can, however, be explained by three main chemical changes that fostered SOM protection.

First, our rock mining residues were clearly pre-weathered (Table 1), therefore, the rocks provided secondary silicate minerals immediately after application that were able to sorb DOM directly. The concentration of carbon within MAOM of these rock-amended soils were lower than their no-rock counterparts, which indicates potential for further carbon sorption and hence MAOM formation with continuing rock weathering.

Secondly, rock provided Ca and Mg that are key cations involved in polyvalent cation bridging and associated MAOM formation, along with Fe and Al. Ca and Mg mostly operate in soil at neutral pH and Fe and Al in acidic conditions (Rowley et al., 2018; Singh et al., 2018). Our baseline soil pH was slightly acidic with a pH of 5.7 in water and 4.9 in 0.01 M CaCl_2_, which increased to 6.2 in water (data not shown) and 5.2 in 0.01 M CaCl_2_ at the end of the incubation across all treatment. At this pH range both groups of cations are similarly important in cation bridging, yet given our correlation analysis (Figure 5), exchangeable Ca and Mg seemed to play a more important role in MAOM formation in our case.

Third, the results from both fractionation assays showed that rock clearly increased SOM occluded within microaggregates. This increase in microaggregates and associated AggOM can be explained by the supply of available Ca and Mg by the rock (Figure 6B), which facilitated the formation of soil aggregates and carbon protection (Baldock, 1989; Clough & Skjemstad, 2000; Rowley et al., 2018; Totsche et al., 2018). Our strong correlation between soil exchangeable Ca and SOM content (Figure 5A) has also been seen in a previous study (Rowley et al., 2018). In our trial, rock addition protected an extra 17% (Figure 6) (or ∼0.15% C in absolute values (Figure 3)) of soil carbon within AggOM that was lost without rock addition. This effect could play a significant role in protection of POM from decomposition, in particular in soil with low degree of aggregation and soil exchangeable Ca and Mg levels.

Overall, the soil carbon content was 32% higher after rock amendment or 0.2% per weight of soil. At a soil bulk density of 1.2 g cm^-3^ and a soil depth of 0.2 m this corresponds to 4.8 t of extra stable carbon stored per hectare. While these results so far are only valid for sandy soils of similar chemistry and cannot be extrapolated to larger areas globally, it does clearly demonstrate the potential of ground rock application for sequestering additional organic carbon. Plants, however, partially counteracted this effect.

### 4.3 Plant counteraction of protection of carbon in aggregates due to micronutrient deficiency

In the absence of rock, plants increased the soil carbon content (Figure 3). However, under the altered chemical conditions after rock-amendment, plants reduced the protection of SOM in aggregates (Figure 6B). It was shown previously that plant root exudates can accelerate the turnover of aggregates, i.e., induce aggregate formation but also destruction (He et al., 2020; Ma et al., 2022; Wang et al., 2020). Plants altered the chemical changes induced by rock addition on SOM content and exchangeable cation levels.

Plants, including wheat, can increase root exudation as a response to nutrient deficiency to solubilise micronutrients, such as Mn, Zn or Fe, to increase their availability and uptake (Awad et al., 1994; Cakmak & Marschner, 1988; Gherardi & Rengel, 2004; F. Li et al., 2018). Zn deficiency in various plant species, for example, increased root exudation by a factor of 2 on average (Cakmak & Marschner, 1988). Such exudates include oxalate, tartarate, L-malate, lactate, citrate and succinate (Gherardi & Rengel, 2004). The dramatic drop in exchangeable micronutrient levels in our study after rock addition, as also observed for Mn in a previous study after basalt application (Anda et al., 2015), and subsequent lower plant uptake suggests exudation to solubilise micronutrients and increase plant uptake (Figure 6B). Availability of Mn and Zn decreases exponentially in the pH range of our soils (5-6.5) through an increase of adsorbed Mn and Zn to soil surfaces (Basta et al., 2005). Rock addition increased the pH by ∼0.2 units within this pH range explaining the drop in micronutrient availability.

Root exudates, such as oxalic acid/oxalate, increase micronutrient availability and hence plant uptake. However, they can also strip polyvalent cations from their metalorganic ligand complexes that results in both loss of cations and carbon (Keiluweit et al., 2015; F. Li et al., 2018; H. Li et al., 2021). Fe does play a particularly important role in the formation of microaggregates (52-250 um) (Z. Lin et al., 2022; Xue et al., 2019) and hence loss of Fe in addition to Ca and Mg can explain the loss in microaggregates and carbon in AggOM in our study. With a loss of Ca, Mg and Fe as mediators between clay surfaces and soil organic carbon and cementation agents, soil aggregation decreased and hence less soil organic carbon was stabilised. This shows the complexity of the system and how soil chemistry changes can alter the effect of plants on SOM content, which in this case resulted in loss of soil carbon with rock addition. Addressing micronutrient deficiencies should avoid the effect plants had on soil aggregation and associated carbon contained within aggregates. Future studies should be designed to specifically investigate this hypothesis.

## Conclusion

We found evidence that a blend of granite and basalt applied to a sandy soil weathered during a 6-month incubation, as demonstrated by soil exchangeable Ca and Mg that were elevated compared to the baseline values. However, this was not associated with the expected increase in soil inorganic carbon content. Instead, rock addition increased SOM stabilisation through the release of Ca and Mg and provision of secondary minerals. A growing wheat plant partially counteracted this affect likely due to the release of plant root exudates induced by reductions in micronutrient levels, Mn and Zn, after rock addition and its associated pH increase. Such exudates solubilised, and hence induced losses of Ca, Mg and Fe that are typically involved in aggregate stabilisation, which also induced losses of carbon formerly protected in aggregates. Still, the application of Ca– and Mg-rich silicates can be a valuable tool to stabilise SOM, particularly in sandy soil and when micronutrient deficiencies are addressed. This could substantially improve the carbon sequestration potential of ground rock application on agricultural land. Higher soil organic carbon levels can have further soil and plant benefits, such as increasing nutrient and water retention. These findings could boost the economic and environmental attractiveness of enhanced rock weathering as a global method for carbon dioxide removal.

## Acknowledgement

The funding for this project was provided by an ANU Grand Challenge. We thank Munash Organics for the provision of the rock for the trials, the Australian Plant Phenomics Facility, which is supported under the National Collaborative Research Infrastructure Strategy of the Australian Government, and Environmental Analysis Laboratory. We acknowledge assistance from Dr. Michael Wellington, Dr. Ulrike Troitzsch, Dr. James Latimer, Dr. Brett Knowles, Robin Grun, Andi Wibowo, Sophia Cain, Imelda Forteza and Alek Meade for lab and technical support.

