## Supplementary Material for "Stabilisation of soil organic matter with rock dust partially counteracted by plants"

Wolfram Buss^a*^

Heath Hasemer^a^

Scott Ferguson^a^

Justin Borevitz^a^

^a^ Research School of Biology, Australian National University, 134 Linnaeus Way, 2601 Canberra, Australia

**Results and Discussion**

**Carbon fractionation assays**

We used two different fractionation assays in this study, a three-pool fractionation technique that keeps soil aggregates intact and hence can distinguish between free POM, AggOM and MAOM, and a two-pool fractionation technique that does not measure POM occluded in aggregates separately but as part of one POM pool that includes free and occluded POM. Both fractionation techniques have advantages and disadvantages. The two-pool fractionation comprises of a harsh extraction technique capable of breaking up POM that subsequently fits through the sieve that separates POM from MAOM. This can mechanically increase the MAOM fraction ^33^. Furthermore, our test shows that using this technique, there is a large DOM pool resulting from the harsh disaggregation step (0.12-0.15% carbon vs. no detectable carbon in the aqueous fraction of the less harsh 3-pool extraction). This DOM fraction is typically not measured ^14,33^, which would underestimate total soil carbon by 10-20% in our study.

The three-pool technique yields an AggOM fraction that contains both occluded POM and MAOM, while the two-pool technique does not consider any physical protection of SOM. There is good evidence that SOM is protected by aggregates on medium-term, which resulted in global promotion of conservation farming techniques, such as no-till ^38–41^. Not considering this AggOM pool is certainly a simplification, and, in this study, we find that rock addition to sandy soil protects SOM in aggregates. The three-pool fractionation requires an extra density separation step with sodium iodide or equivalent high-density solutions, which is more time consuming than simple separation into two pools. However, the high-throughput method described here enables three-pool fractionation of ~50 samples per day by one user, so fractionation of up to 1000 samples per month. Therefore, a three-pool soil carbon fractionation should be the method of choice to ensure both modes of soil carbon stabilisation are considered.

**SI Table 1**: Soil characteristics.

| Parameter | Method reference |  |
| --- | --- | --- |
| Phosphorus (mg/kg P) | Rayment & Lyons 2011 - 9B2 (Colwell) | 31 |
| Nitrate Nitrogen (mg/kg N) | (KCl) | 11 |
| Ammonium Nitrogen (mg/kg N) |  | 14 |
| Sulfur (mg/kg S) |  | 5.2 |
| Exchangeable Calcium (cmol+/kg) | Rayment & Lyons 2011 - 15D3  (Ammonium Acetate) | 0.89 |
| Exchangeable Calcium (mg/kg) |  | 178 |
| Exchangeable Magnesium (cmol+/kg) |  | 0.51 |
| Exchangeable Magnesium (mg/kg) |  | 62 |
| Exchangeable Potassium (cmol+/kg) |  | 0.58 |
| Exchangeable Potassium (mg/kg) |  | 227 |
| Exchangeable Sodium (cmol+/kg) |  | <0.065 |
| Exchangeable Sodium (mg/kg) |  | <15 |
| Exchangeable Aluminium (cmol+/kg) | (KCl) | 0.09 |
| Exchangeable Aluminium (mg/kg) |  | 8.5 |
| Exchangeable Hydrogen (cmol+/kg) | Rayment & Lyons 2011 - 15G1  (Acidity Titration) | 0.22 |
| Exchangeable Hydrogen (mg/kg) |  | 2.2 |
| Effective Cation Exchange Capacity (ECEC) (cmol_+_/kg) | Calculation:  Sum of Ca,Mg,K,Na,Al,H (cmol_+_/kg) | 2.3 |
| Calcium (%) | Base Saturation Calculations -  Cation cmol_+_/kg / ECEC x 100 | 38 |
| Magnesium (%) |  | 22 |
| Potassium (%) |  | 25 |
| Sodium - ESP (%) |  | 1.8 |
| Aluminium (%) |  | 4.1 |
| Hydrogen (%) |  | 9.3 |
| Calcium/Magnesium Ratio | Calculation: Calcium / Magnesium (cmol_+_/kg) | 1.7 |
| Zinc (mg/kg) | Rayment & Lyons 2011 - 12A1 (DTPA) | 1.1 |
| Manganese (mg/kg) |  | 9.2 |
| Iron (mg/kg) |  | 91 |
| Copper (mg/kg) |  | 0.26 |
| Boron (mg/kg) | Rayment & Lyons 2011 - 12C2 (Hot CaCl_2_) | 0.34 |
| Silicon (mg/kg Si) | (Hot CaCl2) | 23 |
| Chloride Estimate (equiv. mg/kg) | Calculation: Electrical Conductivity x 640 | 31 |
| pH in CaCl_2_ | Rayment & Lyons 2011 - 4B4 (CaCl_2_) | 4.9 |
| pH | Rayment & Lyons 2011 - 4A1 (1:5 Water) | 5.68 |
| Electrical Conductivity (dS/m) | Rayment & Lyons 2011 - 3A1 (1:5 Water) | 0.048 |
| Sand (0.05-2 mm) | mastersizer | 84.7 |
| Silt (0.05 - 0.002 mm) | mastersizer | 14.7 |
| Clay (< 0.002 mm) | mastersizer | 0.6 |
| Soil texture | USDA | loamy sand |
| Total nitrogen content | Combustion (%) | 0.045 |
| Total carbon content | Combustion (%) | 0.88 |

**SI Table 2**: Rock elemental content determined via method ME-MS61.

| element | Ag | Al | As | Ba | Be | Bi | Ca | Cd | Ce | Co | Cr | Cs | Cu |
| --- | --- | --- | --- | --- | --- | --- | --- | --- | --- | --- | --- | --- | --- |
| unit | ppm | % | ppm | ppm | ppm | ppm | % | ppm | ppm | ppm | ppm | ppm | ppm |
| content | 0.08 | 6.98 | 1.1 | 360 | 3.44 | 0.48 | 3.22 | 0.1 | 58.5 | 28.5 | 249 | 11.05 | 33.8 |
| element | Fe | Ga | Ge | Hf | In | K | La | Li | Mg | Mn | Mo | Na | Nb |
| unit | % | ppm | ppm | ppm | ppm | % | ppm | ppm | % | ppm | ppm | % | ppm |
| content | 5.48 | 21.8 | 0.17 | 4.5 | 0.073 | 2.19 | 28.2 | 55.4 | 2.29 | 787 | 1.9 | 2.4 | 31 |
| element | Ni | P | Pb | Rb | Re | S | Sb | Sc | Se | Sn | Sr | Ta | Te |
| unit | ppm | ppm | ppm | ppm | ppm | % | ppm | ppm | ppm | ppm | ppm | ppm | ppm |
| content | 94.4 | 1780 | 11.7 | 142.5 | <0.002 | 0.01 | 0.12 | 14.4 | 1 | 6.7 | 404 | 2.68 | <0.05 |
| element | Th | Ti | Tl | U | V | W | Y | Zn | Zr | Hg | Cl | B | Ge |
| unit | ppm | % | ppm | ppm | ppm | ppm | ppm | ppm | ppm | ppm | % | ppm | ppm |
| content | 6.79 | 0.771 | 0.63 | 4.2 | 91 | 3.2 | 20.9 | 100 | 172 | <0.005 | 0.05 | 10 | 2 |

% of particles

| particle size (µm) | soil | basalt |
| --- | --- | --- |
| 0.011482 | 0 | 0 |
| 0.013183 | 0 | 0 |
| 0.015136 | 0 | 0 |
| 0.017378 | 0 | 0 |
| 0.019953 | 0 | 0 |
| 0.022909 | 0 | 0 |
| 0.026303 | 0 | 0 |
| 0.0302 | 0 | 0 |
| 0.034674 | 0 | 0 |
| 0.039811 | 0 | 0 |
| 0.045709 | 0 | 0 |
| 0.052481 | 0 | 0 |
| 0.060256 | 0 | 0 |
| 0.069183 | 0 | 0 |
| 0.079433 | 0 | 0 |
| 0.091201 | 0 | 0 |
| 0.104713 | 0 | 0 |
| 0.120226 | 0 | 0 |
| 0.138038 | 0 | 0 |
| 0.158489 | 0 | 0 |
| 0.18197 | 0 | 0 |
| 0.20893 | 0 | 0 |
| 0.239883 | 0 | 0 |
| 0.275423 | 0 | 0 |
| 0.316228 | 0 | 0 |
| 0.363078 | 0 | 0 |
| 0.416869 | 0 | 0 |
| 0.47863 | 0 | 0 |
| 0.549541 | 0 | 0 |
| 0.630957 | 0 | 0 |
| 0.724436 | 0 | 0 |
| 0.831764 | 0 | 0 |
| 0.954993 | 0 | 0 |
| 1.09648 | 0.02103 | 0.019275 |
| 1.25893 | 0.080409 | 0.501089 |
| 1.44544 | 0.150105 | 1.02673 |
| 1.65959 | 0.18403 | 0.825359 |
| 1.90546 | 0.211586 | 0.90057 |
| 2.18776 | 0.243113 | 1.11821 |
| 2.51189 | 0.281502 | 1.34377 |
| 2.88403 | 0.308405 | 1.56881 |
| 3.31131 | 0.321251 | 1.80056 |
| 3.80189 | 0.331848 | 1.99594 |
| 4.36516 | 0.34343 | 2.07783 |
| 5.01187 | 0.357916 | 2.11244 |
| 5.7544 | 0.379215 | 2.27289 |
| 6.60693 | 0.404168 | 2.60105 |
| 7.58578 | 0.422553 | 2.9399 |
| 8.70964 | 0.428894 | 3.18036 |
| 10 | 0.424776 | 3.346 |
| 11.4815 | 0.42191 | 3.47306 |
| 13.1826 | 0.429704 | 3.59011 |
| 15.1356 | 0.452762 | 3.73617 |
| 17.378 | 0.491027 | 3.92062 |
| 19.9526 | 0.53825 | 4.13348 |
| 22.9087 | 0.580519 | 4.31921 |
| 26.3027 | 0.618292 | 4.4321 |
| 30.1995 | 0.685243 | 4.47513 |
| 34.6737 | 0.852202 | 4.49875 |
| 39.8107 | 1.19335 | 4.55027 |
| 45.7088 | 1.73994 | 4.64258 |
| 52.4807 | 2.4283 | 4.73345 |
| 60.256 | 3.09286 | 4.71616 |
| 69.1831 | 3.54459 | 4.48255 |
| 79.4328 | 3.67676 | 3.97314 |
| 91.2011 | 3.53274 | 3.17507 |
| 104.713 | 3.26282 | 2.14882 |
| 120.226 | 3.04768 | 1.10913 |
| 138.038 | 3.07186 | 0.259416 |
| 158.489 | 3.50058 | 0 |
| 181.97 | 4.40667 | 0 |
| 208.93 | 5.71949 | 0 |
| 239.883 | 7.05278 | 0 |
| 275.423 | 7.96752 | 0 |
| 316.228 | 8.10447 | 0 |
| 363.078 | 7.36184 | 0 |
| 416.869 | 5.97407 | 0 |
| 478.63 | 4.42106 | 0 |
| 549.541 | 3.06848 | 0 |
| 630.957 | 2.04469 | 0 |
| 724.436 | 1.21962 | 0 |
| 831.764 | 0.460799 | 0 |
| 954.993 | 0.142851 | 0 |
| 1096.48 | 0 | 0 |

% of particles

| particle size (µm) | soil | basalt |
| --- | --- | --- |
| 0.011482 | 0 | 0 |
| 0.013183 | 0 | 0 |
| 0.015136 | 0 | 0 |
| 0.017378 | 0 | 0 |
| 0.019953 | 0 | 0 |
| 0.022909 | 0 | 0 |
| 0.026303 | 0 | 0 |
| 0.0302 | 0 | 0 |
| 0.034674 | 0 | 0 |
| 0.039811 | 0 | 0 |
| 0.045709 | 0 | 0 |
| 0.052481 | 0 | 0 |
| 0.060256 | 0 | 0 |
| 0.069183 | 0 | 0 |
| 0.079433 | 0 | 0 |
| 0.091201 | 0 | 0 |
| 0.104713 | 0 | 0 |
| 0.120226 | 0 | 0 |
| 0.138038 | 0 | 0 |
| 0.158489 | 0 | 0 |
| 0.18197 | 0 | 0 |
| 0.20893 | 0 | 0 |
| 0.239883 | 0 | 0 |
| 0.275423 | 0 | 0 |
| 0.316228 | 0 | 0 |
| 0.363078 | 0 | 0 |
| 0.416869 | 0 | 0 |
| 0.47863 | 0 | 0 |
| 0.549541 | 0 | 0 |
| 0.630957 | 0 | 0 |
| 0.724436 | 0 | 0 |
| 0.831764 | 0 | 0 |
| 0.954993 | 0 | 0 |
| 1.09648 | 0.02103 | 0.019275 |
| 1.25893 | 0.101439 | 0.520364 |
| 1.44544 | 0.251544 | 1.5471 |
| 1.65959 | 0.435574 | 2.37245 |
| 1.90546 | 0.64716 | 3.27303 |
| 2.18776 | 0.890273 | 4.39123 |
| 2.51189 | 1.17178 | 5.73501 |
| 2.88403 | 1.48018 | 7.30381 |
| 3.31131 | 1.80143 | 9.10438 |
| 3.80189 | 2.13328 | 11.1003 |
| 4.36516 | 2.47671 | 13.1781 |
| 5.01187 | 2.83462 | 15.2906 |
| 5.7544 | 3.21384 | 17.5635 |
| 6.60693 | 3.61801 | 20.1645 |
| 7.58578 | 4.04056 | 23.1044 |
| 8.70964 | 4.46945 | 26.2848 |
| 10 | 4.89423 | 29.6308 |
| 11.4815 | 5.31614 | 33.1038 |
| 13.1826 | 5.74584 | 36.694 |
| 15.1356 | 6.19861 | 40.4301 |
| 17.378 | 6.68963 | 44.3507 |
| 19.9526 | 7.22788 | 48.4842 |
| 22.9087 | 7.8084 | 52.8034 |
| 26.3027 | 8.42669 | 57.2355 |
| 30.1995 | 9.11194 | 61.7107 |
| 34.6737 | 9.96414 | 66.2094 |
| 39.8107 | 11.1575 | 70.7597 |
| 45.7088 | 12.8974 | 75.4023 |
| 52.4807 | 15.3257 | 80.1357 |
| 60.256 | 18.4186 | 84.8519 |
| 69.1831 | 21.9632 | 89.3344 |
| 79.4328 | 25.64 | 93.3076 |
| 91.2011 | 29.1727 | 96.4826 |
| 104.713 | 32.4355 | 98.6315 |
| 120.226 | 35.4832 | 99.7406 |
| 138.038 | 38.5551 | 100 |
| 158.489 | 42.0556 | 100 |
| 181.97 | 46.4623 | 100 |
| 208.93 | 52.1818 | 100 |
| 239.883 | 59.2346 | 100 |
| 275.423 | 67.2021 | 100 |
| 316.228 | 75.3066 | 100 |
| 363.078 | 82.6684 | 100 |
| 416.869 | 88.6425 | 100 |
| 478.63 | 93.0636 | 100 |
| 549.541 | 96.132 | 100 |
| 630.957 | 98.1767 | 100 |
| 724.436 | 99.3963 | 100 |
| 831.764 | 99.8571 | 100 |
| 954.993 | 100 | 100 |
| 1096.48 | 100 | 100 |
| 1258.93 | 100 | 100 |
| 1445.44 | 100 | 100 |
| 1659.59 | 100 | 100 |
| 1905.46 | 100 | 100 |
| 2187.76 | 100 | 100 |

**SI Figure 1**: Size distribution of particles in soil and rock per (A) size class and (B) accumulated.

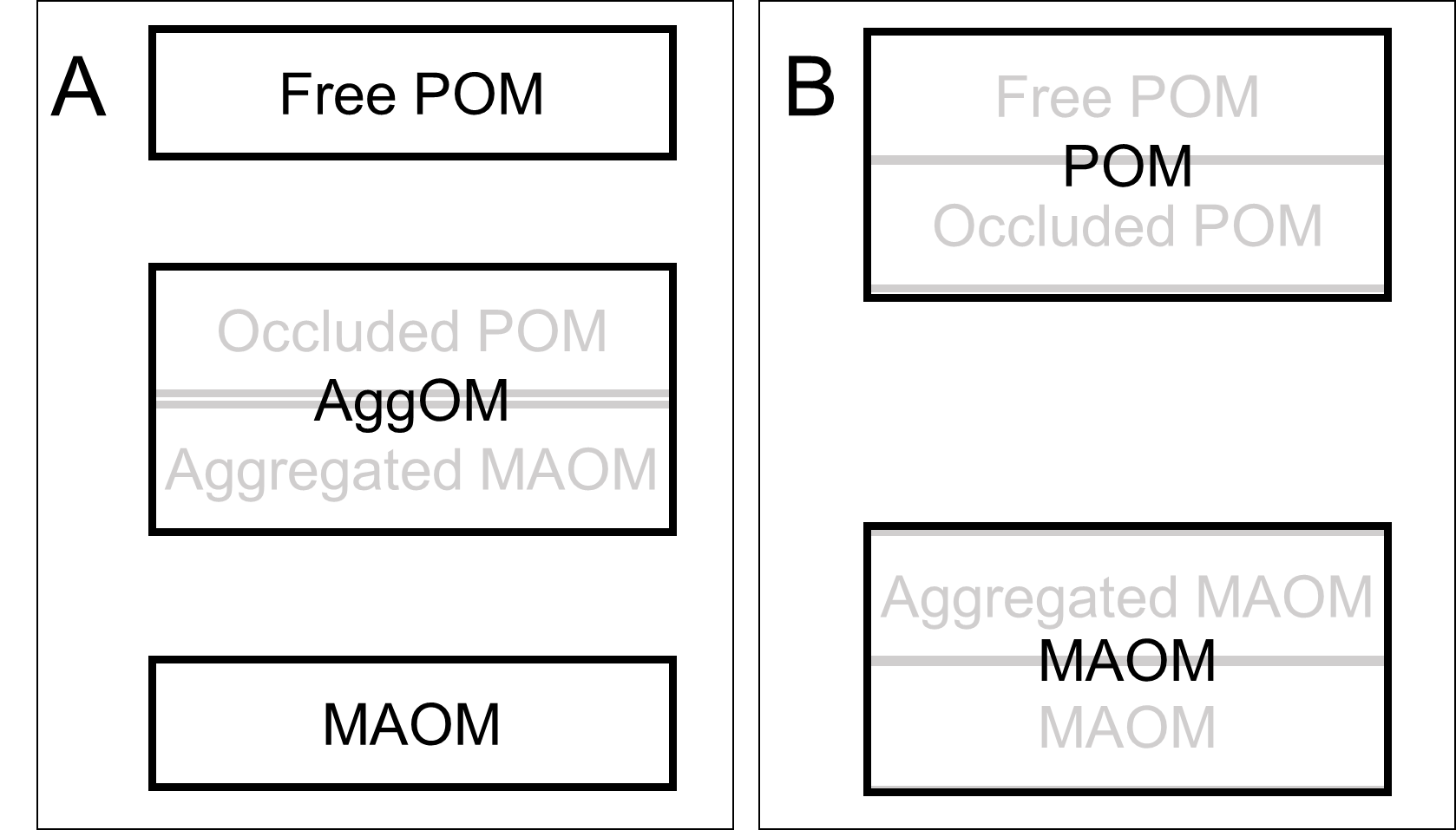

**SI Figure 2**: Schematic representation of fractions in (A) three-pool and (B) two-pool soil carbon fractionation assays.

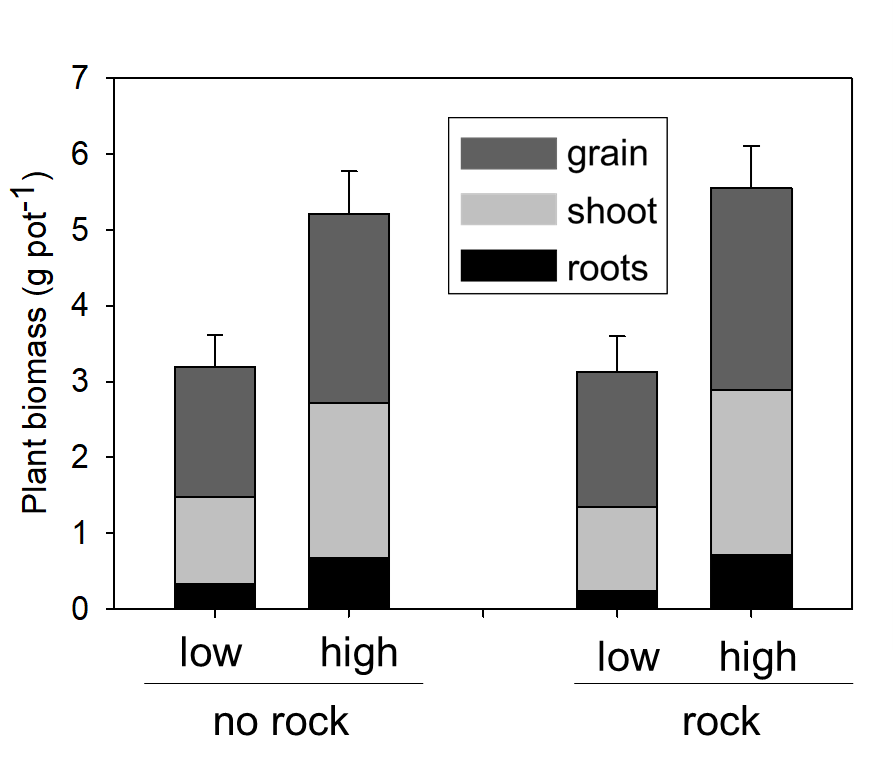

**SI Figure 3**: Plant (wheat) biomass separated into grain, shoot and root as affected by rock addition and high/low water treatment at the end of the 6-month trial.

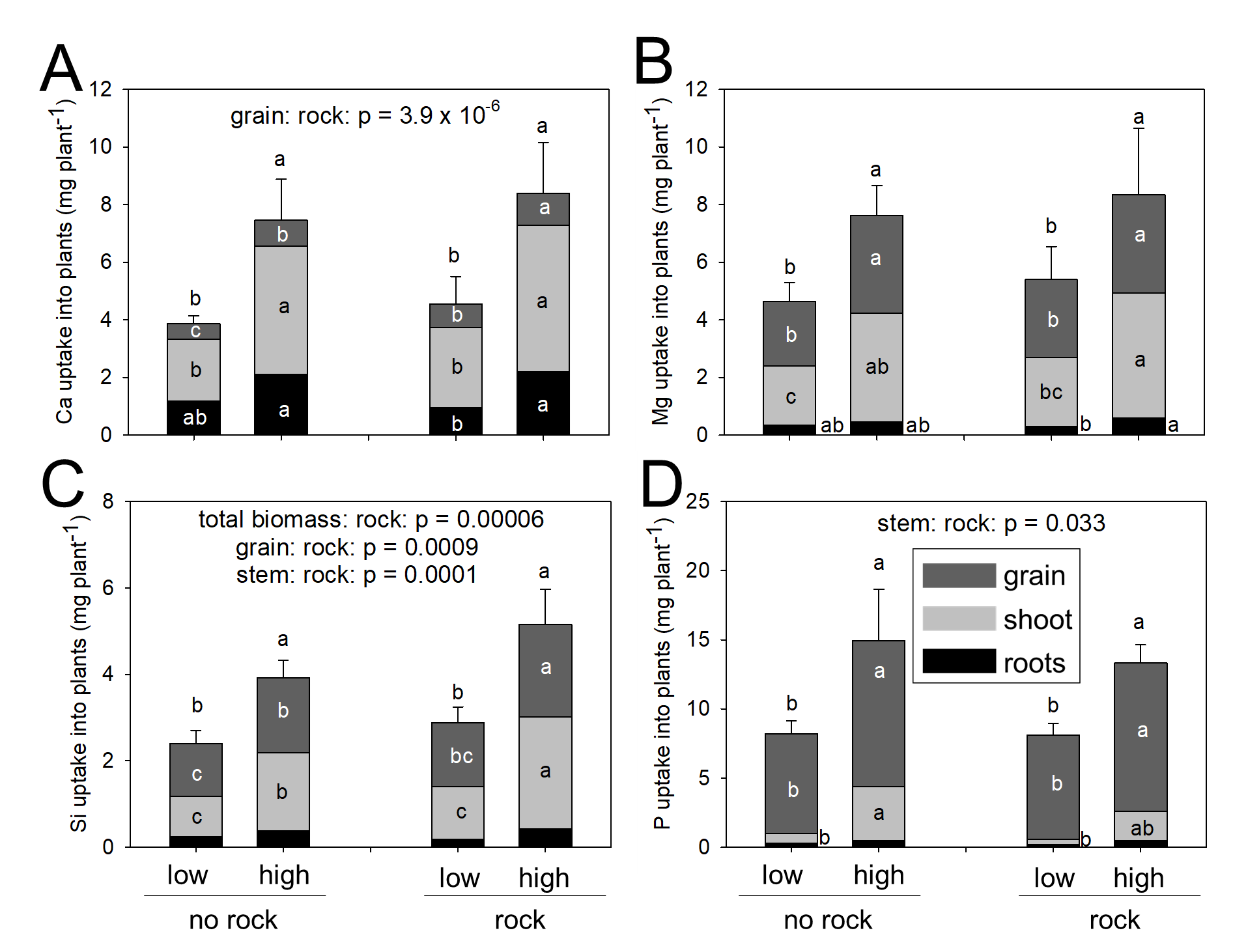

**SI Figure 4**: Uptake of Ca, Mg, Si and P in different plant tissue (root, shoot and grain) at the end of a 6-month incubation. Main effects determined by one-way ANOVAs shown above each figure. Different letters indicate significant differences among the treatments determined via Tukey post-hoc test. Water had a significant effect on all elemental plant tissue contents (mg plant^-1^).

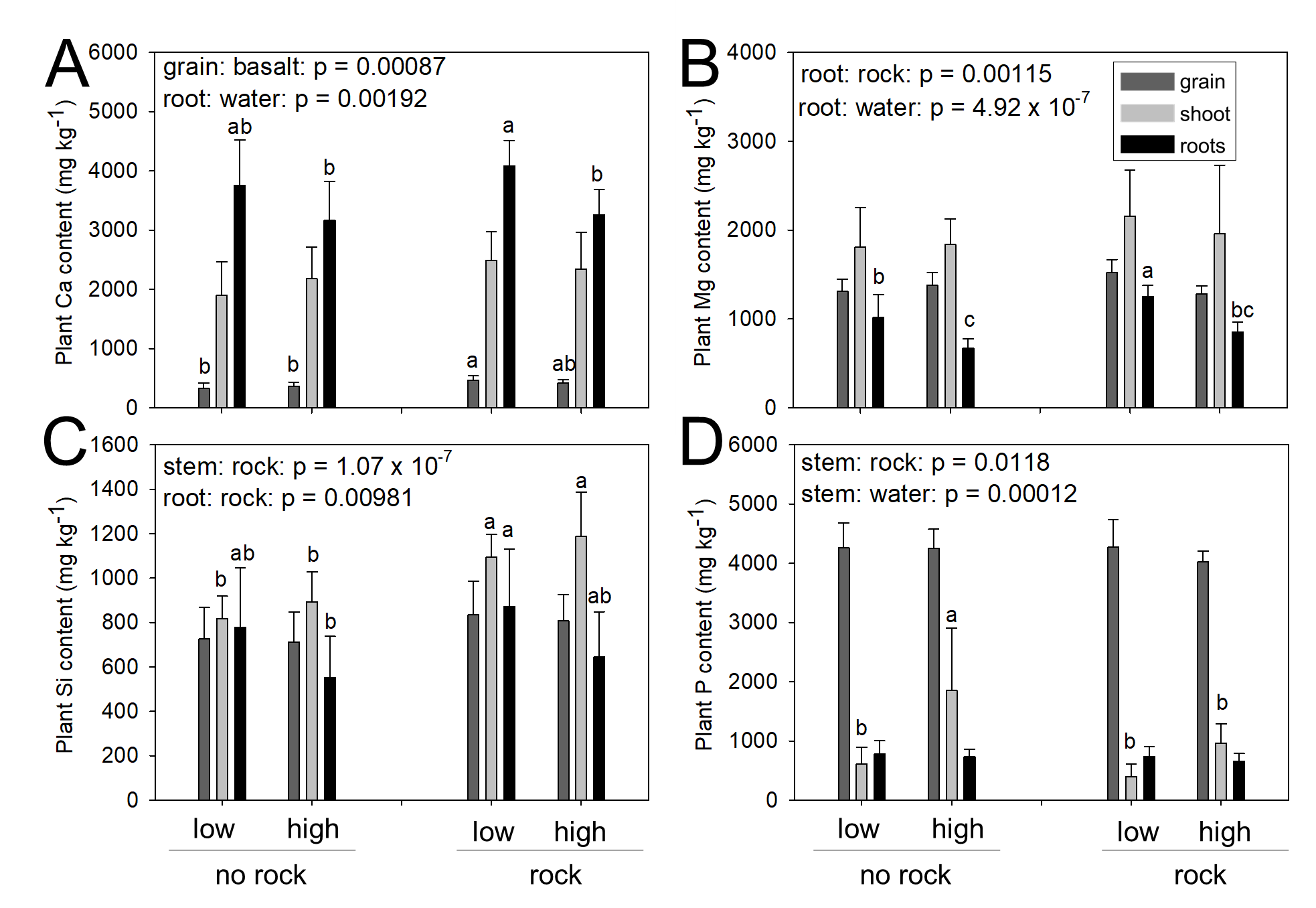

**SI Figure 5**: Plant tissue contents of (A) Ca, (B) Mg, (C) Si and (D) P in grain, stem and root at the end of a 6-months incubation. Main effects determine by one-way ANOVAs shown above each figure. Different letters indicate significant differences among the treatments determined via Tukey post-hoc test.

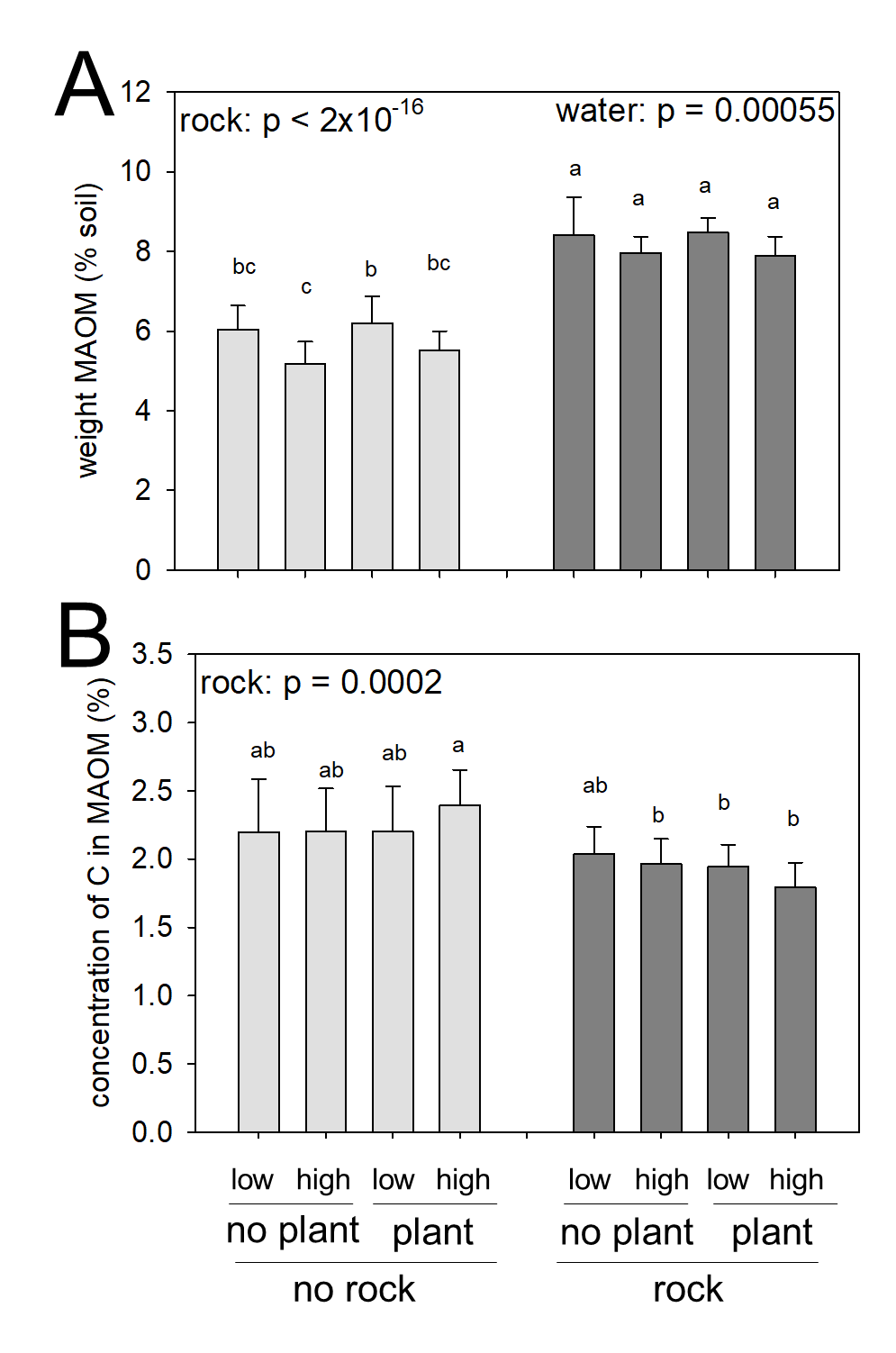

**SI Figure 6**: Amount of soil as mineral-associated organic matter at the end of a 6-months incubation. Full factorial design testing high vs. low water, unplanted vs. planted and soil only vs. rock addition.

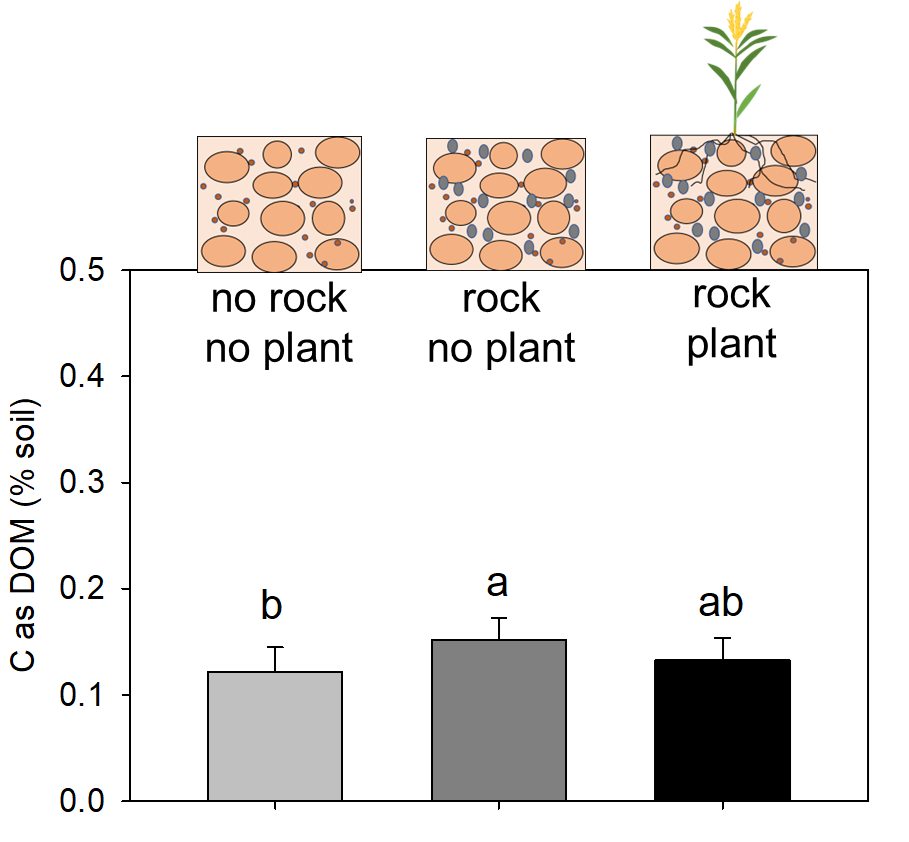

**SI Figure 7**: Carbon content present as dissolved organic matter (DOM) in the aqueous fraction of the two-pool soil carbon fractionation. Treatments: no rock without plant (control-control-low), rock with and without plants (all low water treatments). Soil carbon fractionation using hexameta-phosphate extraction for full soil disaggregation at the end of a 6-months incubation. Different letters indicate significant differences among the treatments determined via Tukey post-hoc test.

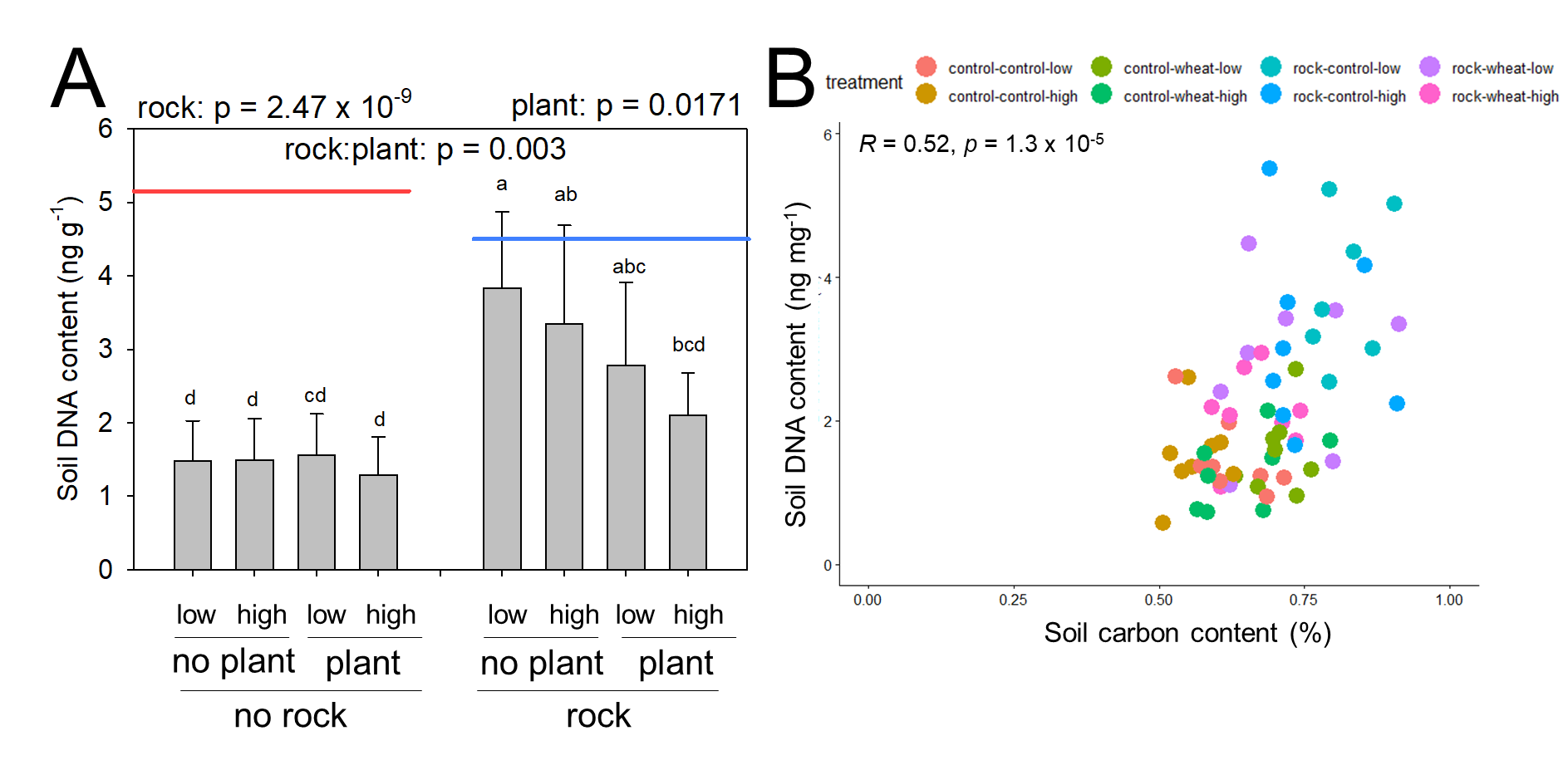

**SI Figure 8**: Relationship of soil DNA content and carbon content at the end of a 6-months incubation. Full factorial design testing high vs. low water, unplanted vs. planted and soil only vs. rock addition. Red and blue lines indicate baseline values from the soil at the start of the trial without and with rock addition, respectively.

**SI Table 3**: Read length, total reads and total base pairs of soil DNA sequencing.

| ID | min read length | max read length | total reads | total base pairs | average read length |
| --- | --- | --- | --- | --- | --- |
| control-control-low-1 | 200 | 15539 | 21359 | 13900541 | 651 |
| control-control-low-2 | 200 | 14061 | 14498 | 18538858 | 1279 |
| control-control-low-3 | 200 | 14222 | 29963 | 53246392 | 1777 |
| control-control-low-4 | 200 | 18429 | 28888 | 54167880 | 1875 |
| control-control-low-5 | 200 | 19931 | 48279 | 73441780 | 1521 |
| control-control-low-6 | 200 | 12405 | 18719 | 29115956 | 1555 |
| control-control-low-7 | 200 | 26388 | 8095 | 21140003 | 2611 |
| control-control-low-8 | 200 | 20239 | 23591 | 48380423 | 2051 |
| control-control-high-1 | 200 | 216077 | 21057 | 14365872 | 682 |
| control-control-high-2 | 200 | 13698 | 12873 | 17785610 | 1382 |
| control-control-high-3 | 200 | 11564 | 12082 | 17450634 | 1444 |
| control-control-high-4 | 200 | 17820 | 8380 | 12131525 | 1448 |
| control-control-high-5 | 200 | 18863 | 38397 | 48549951 | 1264 |
| control-control-high-6 | 200 | 22395 | 10119 | 16202747 | 1601 |
| control-control-high-7 | 200 | 43127 | 5789 | 18019649 | 3113 |
| control-control-high-8 | 200 | 10338 | 6647 | 8342228 | 1255 |
| control-wheat-low-1 | 200 | 15754 | 17210 | 11410405 | 663 |
| control-wheat-low-2 | 200 | 14789 | 25749 | 38210410 | 1484 |
| control-wheat-low-3 | 200 | 9424 | 4907 | 7971484 | 1625 |
| control-wheat-low-4 | 200 | 14929 | 15040 | 25651172 | 1706 |
| control-wheat-low-5 | 200 | 31149 | 36809 | 51316293 | 1394 |
| control-wheat-low-6 | 200 | 22623 | 21972 | 29705547 | 1352 |
| control-wheat-low-7 | 200 | 11775 | 7658 | 11158993 | 1457 |
| control-wheat-low-8 | 200 | 32161 | 10239 | 24370568 | 2380 |
| control-wheat-high-1 | 200 | 9038 | 40407 | 26481411 | 655 |
| control-wheat-high-2 | 200 | 17002 | 17815 | 28238332 | 1585 |
| control-wheat-high-3 |  |  |  |  |  |
| control-wheat-high-4 | 200 | 18660 | 17299 | 36308761 | 2099 |
| control-wheat-high-5 | 200 | 68951 | 9158 | 12787669 | 1396 |
| control-wheat-high-6 | 200 | 12396 | 17787 | 24401121 | 1372 |
| control-wheat-high-7 | 201 | 141206 | 9472 | 26188898 | 2765 |
| control-wheat-high-8 | 201 | 12032 | 4517 | 6490494 | 1437 |
| rock-control-low-1 | 200 | 9746 | 8150 | 11445262 | 1404 |
| rock-control-low-2 | 200 | 17009 | 14241 | 18128464 | 1273 |
| rock-control-low-3 | 200 | 10599 | 11252 | 17711609 | 1574 |
| rock-control-low-4 | 200 | 45451 | 37372 | 74583512 | 1996 |
| rock-control-low-5 | 200 | 26304 | 28742 | 44407495 | 1545 |
| rock-control-low-6 | 200 | 11974 | 10316 | 16658427 | 1615 |
| rock-control-low-7 | 200 | 30844 | 12999 | 41992251 | 3230 |
| rock-control-low-8 | 200 | 24349 | 18046 | 42998031 | 2383 |
| rock-control-high-1 | 200 | 13412 | 23218 | 16497707 | 711 |
| rock-control-high-2 | 200 | 13247 | 18258 | 26928294 | 1475 |
| rock-control-high-3 | 200 | 13792 | 16194 | 29981829 | 1851 |
| rock-control-high-4 | 200 | 20174 | 12883 | 27617167 | 2144 |
| rock-control-high-5 | 200 | 25562 | 25754 | 44972194 | 1746 |
| rock-control-high-6 | 200 | 17123 | 13323 | 19430997 | 1458 |
| rock-control-high-7 | 200 | 25766 | 9314 | 25139807 | 2699 |
| rock-control-high-8 | 200 | 26147 | 8393 | 16055570 | 1913 |
| rock-wheat-low-1 | 200 | 36139 | 40922 | 24997652 | 611 |
| rock-wheat-low-2 | 200 | 11224 | 9585 | 15492384 | 1616 |
| rock-wheat-low-3 | 200 | 27695 | 28430 | 50133412 | 1763 |
| rock-wheat-low-4 | 200 | 19425 | 12185 | 26275436 | 2156 |
| rock-wheat-low-5 | 200 | 117427 | 41798 | 64016813 | 1532 |
| rock-wheat-low-6 | 200 | 38824 | 14167 | 26713152 | 1886 |
| rock-wheat-low-7 | 200 | 36233 | 21088 | 71419494 | 3387 |
| rock-wheat-low-8 | 200 | 11393 | 6923 | 9551596 | 1380 |
| rock-wheat-high-1 | 200 | 14861 | 23498 | 16325935 | 695 |
| rock-wheat-high-2 | 200 | 21257 | 17287 | 23270177 | 1346 |
| rock-wheat-high-3 | 200 | 17273 | 5978 | 11335041 | 1896 |
| rock-wheat-high-4 | 200 | 23525 | 18243 | 36433757 | 1997 |
| rock-wheat-high-5 |  |  |  |  |  |
| rock-wheat-high-6 | 200 | 17579 | 15175 | 27123698 | 1787 |
| rock-wheat-high-7 | 200 | 12734 | 5546 | 8758124 | 1579 |
| rock-wheat-high-8 | 200 | 10084 | 9589 | 12528998 | 1306 |

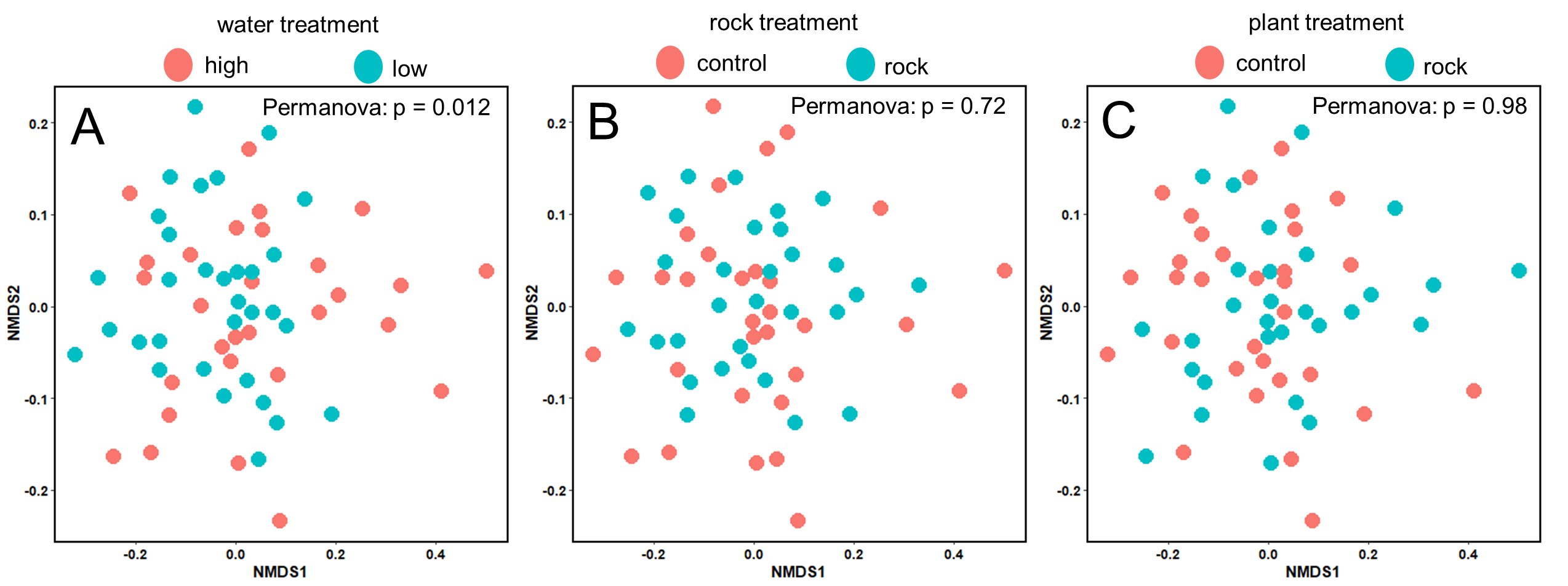

**SI Figure 9**: Effect of (A) water, (B) rock and (C) plant on microbial composition in soil at the end of a 6-months incubation depicted via NMDS.

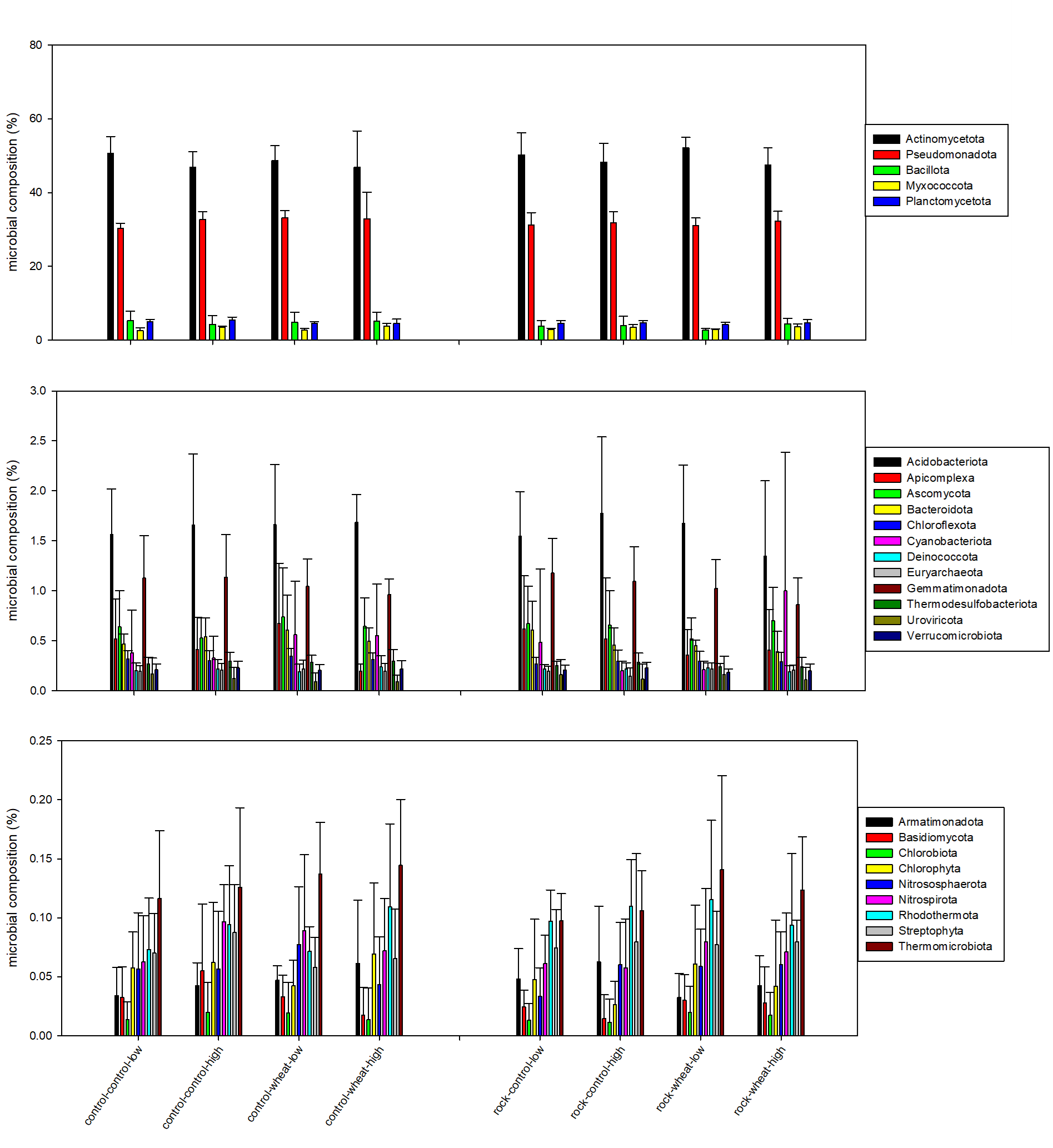

**SI Figure 10:** Phylum level microbial composition (%) in soil at the end of a 6-months incubation. (A) most common phyla (>4%), (B) less common phyla (>0.2%) and (C) least common phyla (<0.2%).

**SI Table 4**: Analysis of compositions of microbiomes with bias correction (ANCOM-BC) with subsequent testing for significant changes using the ANCOM-BC2 package in R on phyla, class and order level comparing basalt, water and plant effects. Only operational taxonomic units with significant changes shown.

|  | Phylum | Class | Order |
| --- | --- | --- | --- |
| basalt | N/A | N/A | N/A |
| plant | N/A | N/A | N/A |
| water | N/A | Rubrobacteria: p = 0.014 | Catenulisporales: p = 0.008 |
|  |  |  | Nostocales: p = 0.018 |
|  |  |  | Rubrobacterales: p = 0.011 |
|  |  |  | Solirubrobacterales: p = 0.040 |

**SI Table 5**: PERMANOVA results for testing differences between basalt, plant and water treatment on phyla, class and order level.

|  | Phylum | Class | Order |
| --- | --- | --- | --- |
| basalt | 0.745 | 0.692 | 0.664 |
| plant | 0.998 | 0.841 | 0.711 |
| water | **0.013** | **0.011** | **0.008** |
